# Temporal variabilities provide additional category-related information in object category decoding: a systematic comparison of informative EEG features

**DOI:** 10.1101/2020.09.02.279042

**Authors:** Hamid Karimi-Rouzbahani, Mozhgan Shahmohammadi, Ehsan Vahab, Saeed Setayeshi, Thomas Carlson

## Abstract

How does the human brain encode visual object categories? Our understanding of this has advanced substantially with the development of multivariate decoding analyses. However, conventional electroencephalography (EEG) decoding predominantly use the “mean” neural activation within the analysis window to extract category information. Such temporal averaging overlooks the within-trial neural variability which is suggested to provide an additional channel for the encoding of information about the complexity and uncertainty of the sensory input. The richness of temporal variabilities, however, has not been systematically compared with the conventional “mean” activity. Here we compare the information content of 31 variability-sensitive features against the “mean” of activity, using three independent highly-varied datasets. In whole-trial decoding, the classical event-related potential (ERP) components of “P2a” and “P2b” provided information comparable to those provided by “Original Magnitude Data (OMD)” and “Wavelet Coefficients (WC)”, the two most informative variability-sensitive features. In time-resolved decoding, the “OMD” and “WC” outperformed all the other features (including “mean”), which were sensitive to limited and specific aspects of temporal variabilities, such as their phase or frequency. The information was more pronounced in Theta frequency band, previously suggested to support feed-forward visual processing. We concluded that the brain might encode the information in multiple aspects of neural variabilities simultaneously e.g. phase, amplitude and frequency rather than “mean” per se. In our active categorization dataset, we found that more effective decoding of the neural codes corresponds to better prediction of behavioral performance. Therefore, the incorporation of temporal variabilities in time-resolved decoding can provide additional category information and improved prediction of behavior.

## Introduction

How does the brain encode information about visual object categories? This question has been studied for decades using different neural recording techniques including invasive neurophysiology (Hung et al., 2005) and electrocorticography (ECoG; Majima et al., 2014; Watrous et al., 2015; Rupp et al., 2017; Lie et al., 2009; Miyakawa et al., 2018; Liu et al., 2009), as well as non-invasive neuroimaging methods such as functional Magnetic Resonance Imaging (fMRI; Haxby et al., 2001), magnetoencephalography (MEG; Contini et al., 2017; Carlson et al., 2013) and electroencephalography (EEG; Kaneshiro et al., 2015; Simanova et al., 2010) or a combination of them (Cichy et al., 2014). There has been great success in “reading-out” or “decoding” neural representations of semantic object categories from neuroimaging data. However, it is still unclear if the conventional decoding analyses effectively detect the complex neural codes. Critically, one potential source of neural codes, in high-temporal-resolution data (e.g. EEG), can be the “within-trial/window temporal variability” of EEG signals, which is generally ignored through temporal averaging in decoding. The use of such summarized “mean” activity, can hide the true spatiotemporal dynamics of neural processes such as object category encoding, which is still debated in cognitive neuroscience (Grootswagers et al., 2019; Majima et al., 2014; Karimi-Rouzbahani et al., 2017b; Isik et al., 2013; Cichy et al., 2014). Here, we quantitatively compare the information content and the temporal dynamics of a large set of features from EEG time series, each sensitive to a specific aspect of within-trial temporal variability. We then evaluate the relevance of these features by measuring how well each one predicts behavioral performance.

Sensory neural codes are multiplexed structures containing information on different time scales and about different aspects of the sensory input (Panzeri et al., 2010; Wark et al., 2009; Gawne et al., 1996). Previous animal studies have shown that the brain does not only encode the sensory information in the neural firing rates (i.e. average number of neural spikes within specific time windows), but also in more complex patterns of neural activity such as millisecond-precise activity and phase (Kayser et al., 2009; Victor, 2000; Montemurro et al., 2008). It was shown that stimulus contrast was represented by latency coding at a temporal precision of ∼10 ms, whereas the stimulus orientation and the spatial frequency were encoded at a coarser temporal precision (30 ms and 100 ms, respectively; Victor, 2000). It was shown that spike rates on 5-10-ms timescales carried complementary information to the phase of firing relative to low-frequency (1-8 Hz) LFPs about epoch of naturalistic movie (Montemurro et al., 2008). Therefore, the temporal patterns/variabilities of neural activity are enriched platforms of neural codes.

Recent computational and experimental studies have proposed that neural variability, provides a separate and additional channel to the “mean” activity, for the encoding of general aspects of the sensory information e.g. its “*uncertainty*” and “*complexity*” (Orbán et al., 2016; Garrett et al., 2020). Specifically, *uncertainty* about the stimulus features (e.g. orientations of lines in the image) was directly linked to neural variability in monkeys’ visual area (Orbán et al., 2016) and human EEG (Kosciessa et al., 2021): wider inferred range of possible feature combinations in the input stimulus corresponded to wider distribution of neural responses. This could be applied to both within- and across-trial variability (Orbán et al., 2016). Moreover, temporal variability was directly related to the *complexity* of input images: higher neural variability for house (i.e. more varied) vs. face (i.e. less varied) images (Garrett et al., 2020) and provided a reliable measure of perceptual performance in behavior (Waschke et al., 2019). The *uncertainty*- and *complexity*-dependent modulation of neural variability, which is linked to the category of input information, has been suggested to facilitate neural energy saving, adaptive and effective encoding of the sensory inputs in changing environments (Garrett et al., 2020; Waschke et al., 2021).

Despite the richness of information encoded by neural variabilities, the unclear transformation of such neuronal codes into EEG activity has led to divergent approaches used for decoding information from EEG. For example, the information in neural firing rates might appear in phase patterns rather than amplitude of EEG oscillations (Ng et al., 2013). Generally, three families of features have been extracted from EEG time series to detect neural codes from temporal variabilities (Waschke et al., 2021): variance-, frequency- and information theory-based features, each detecting specific aspects of variability. In whole-trial decoding, components of event-related potentials (ERPs) such as N1, P1, P2a and P2b, which quantify time-specific variabilities of within-trial activation, have provided significant information about object categories (separately and in combination; Chan et al., 2011; Wang et al., 2012; Qin et al., 2016). Others successfully decoded information from more complex variance- and frequency-based features such as signal phase (Behroozi et al., 2016; Watrous et al., 2015; Torabi et al., 2017; Wang et al., 2018; Voloh et al., 2020), signal power across frequency bands (Rupp et al., 2017; Miyakawa et al., 2018; Majima et al., 2014; Miyakawa et al., 2018), time-frequency Wavelet coefficients (Hatamimajoumerd and Talebpour, 2019; Taghizadeh-Sarabi et al., 2015), inter-electrode temporal correlations (Karimi-Rouzbahani et al., 2017a) and information-based features (e.g. entropy; Joshi et al., 2018; Torabi et al., 2017; Stam, 2005). Therefore, the neural codes are generally detected from EEG activity using a wide range of features sensitive to temporal variability.

While insightful, previous studies have also posed new questions about the relative richness, temporal dynamics and the behavioral relevance of different features of neural variability. First, can the features sensitive to temporal variabilities, provide additional category information to the conventional “mean” feature? While several of the above studies have compared multiple features (Chan et al., 2011; Taghizadeh-Sarabi et al., 2015; Torabi et al., 2016), none of them compared their results against the conventional “mean” activity, which is the dominant feature, especially in time-resolved decoding (Grootswagers et al., 2017). This comparison will not only validate the richness of each feature of neural variability but will also show if the mean activity detects a large portion of the neural codes produced by the brain. We predicted that the informative neural variabilities, if properly decoded, should provide additional information to the “mean” activity, which overlooks the temporal variability within the analysis window.

Second, do the features sensitive to temporal variabilities evolve over similar time windows to the “mean” feature? Among all the studies mentioned above, only a few investigated the temporal dynamics of features, other than the “mean” in ***time-resolved*** decoding (Majima et al., 2014; Stewart et al., 2014; Karimi-Rouzbahani et al., 2017a), where the temporal evolution of information encoding is studied (Grootswagers et al., 2017). As distinct aspects of sensory information (e.g. contrast vs. spatial frequency) are represented on different temporal scales (Victor, 2000; Montemurro et al., 2008) and different variability features are potentially sensitive to distinct aspects of variability, we might see differential temporal dynamics for different features.

Third, do the features sensitive to temporal variabilities explain the behavioral recognition performance more accurately than the “mean” feature? One important question, which was not covered in the above studies, was whether the extracted information was behaviorally relevant or was it just epiphenomenal to the experimental conditions. One way of validating the relevance of the extracted neural codes is to check if they could predict the relevant behavior (Williams et al., 2007; Grootswagers et al., 2018; Woolgar et al., 2019). We previously found that the decoding accuracies obtained from “mean” signal activations could predict the behavioral recognition performance (Ritchie, et al., 2015). However, it remains unknown whether (if at all) the information obtained from temporal variabilities can explain more variance of the behavioral performance. Our prediction was that, as the more informative features access more of the potentially overlooked neural codes, they should also explain the behavioral performance more accurately.

In this study, we address the above questions, to provide additional insights about what aspects of neural variabilities might reflect the neural codes more thoroughly and how we can extract them most effectively using multivariate decoding analyses.

## Methods

The datasets used in this study and the code are available online at https://osf.io/wbvpn/. All the open-source scripts used in this study were compared against other implementations of identical algorithms in simulations and used only if they produced identical results. All open-source implementation scripts of similar algorithms produced identical results in our simulations. To evaluate different implementations, we tested them using 1000 random (normally distributed with unit variance and zero mean) time series each including 1000 samples.

### Overview of datasets

We chose three previously published EEG datasets in this study, which differed across a wide range of parameters including the recording set-up (e.g. amplifier, number of electrodes, preprocessing steps, etc.), characteristics of the image-set (e.g. number of categories and exemplars within each category, colorfulness of images, etc.), and task (e.g. presentation length, order and the participants’ task; Table 1). All three datasets previously successfully provided object category information using multivariate analyses.

**Table 1.** Details of the three datasets used in this study.

| Dataset | # electrodes | Band-pass filtering | Notch filtering | # object categories | # stimulus repetition | Stimulus presentation time | Stimulus size (periphery) | Task | Participants' accuracy | Participants' Age (median) | Participants' gender |
| --- | --- | --- | --- | --- | --- | --- | --- | --- | --- | --- | --- |
| **1** Karimi-Rouzbahani et al., 2017a | 31 | 0.03-200 Hz | 50 Hz | 4 | 3 | 50 ms | 2~13.5° (0.7~8.8°) | Color matching (passive) | %94.68 | 22.1 | 7 male<br>3 female |
| **2** Karimi-Rouzbahani et al., 2019 | 31 | 0.03-200 Hz | 50 Hz | 4 | 6 | 900 ms | 8° × 8° (0) | Object category detection (active) | %94.65 | 26.4 | 6 male<br>4 female |
| **3** Kaneshiro et al., 2015 | 128 | 0.03-50 Hz | No | 6 | 12 | 500 ms | 7.0° × 6.5° (0) | No task (fixation) | N/A | 30.5 | 7 male<br>3 female |

#### Dataset 1

We have previously collected Dataset 1 while participants were briefly (i.e. 50 ms) presented with gray-scale images from four synthetically-generated 3D object categories (Karimi-Rouzbahani et al., 2017a). The objects underwent systematic variations in scale, positional periphery, in-depth rotation and lighting conditions, which made perception difficult, especially in extreme variation conditions. Randomly ordered stimuli were presented in consecutive pairs (Figure 1, top row). The participant’s task was unrelated to object categorization; they pressed one of two pre-determined buttons to indicate if the fixation dots, superimposed on the first and second stimuli, were the same/different color (2-alternative forced choice).

**Figure 1.**
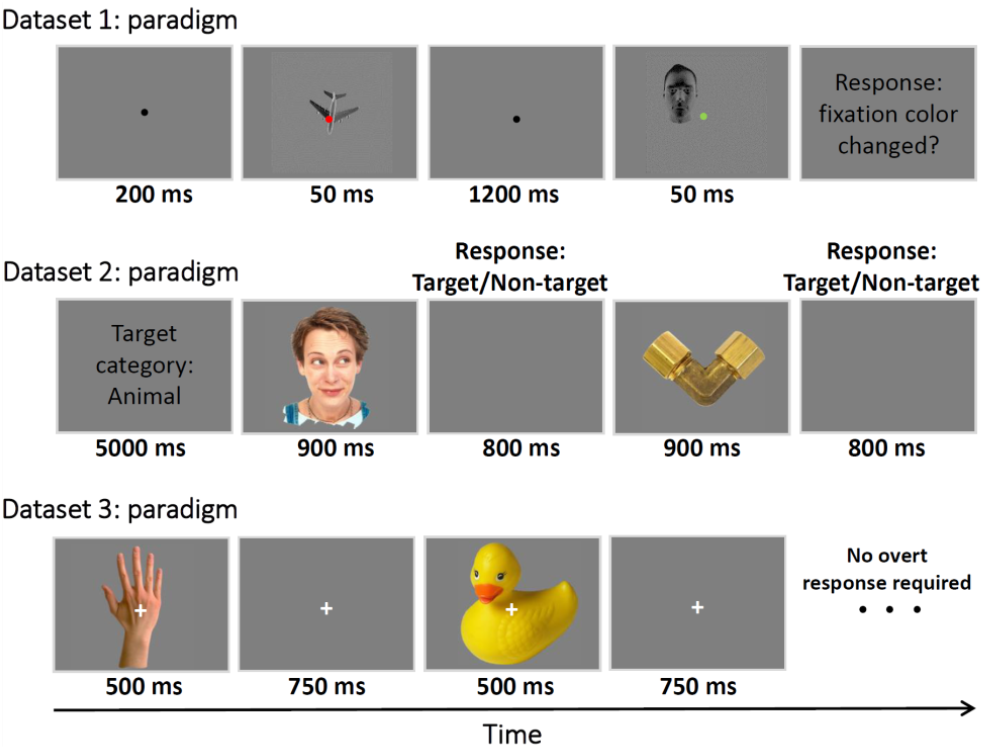
Paradigms of the datasets used in this study. Dataset 1 (top row) presented two consecutive object images each with a fixation dot. Participants’ task was to indicate if the fixation dot was the same or different colors across the image pairs (passive task). Dataset 2 (middle row) presented objects from the target and non-target categories in sequences of 12 images. Participant’s task was to indicate, for each image, if it was from the target/non-target category (active task). Dataset 3 (bottom row), presented sequences of object images from 6 different categories. Participants did not have any specific tasks, except for looking at the center of the image (no overt task). See more details about the datasets in the relevant references provided in Table 1.

#### Dataset 2

We have collected Dataset 2 in an active categorization experiment, in which participants pressed a button if the presented object image was from a target category (go/no-go), which was cued at the beginning of each block of 12 stimuli (Karimi-Rouzbahani et al., 2019; Figure 1, middle row). The object images, which were cropped from real photographs, were part of the well-stablished benchmark image set for object recognition developed by Kiani et al., (2007). This image set has been previously used to extract object category information from both human and monkey brain using MEG (Cichy et al., 2014), fMRI (Cichy et al., 2014; Kriegeskorte et al., 2008) and single-cell electrophysiology (Kriegeskorte et al., 2008; Kiani et al., 2007).

#### Dataset 3

We also used another dataset (Dataset 3) which was not collected in our lab. This dataset was collected by Kaneshiro et al., (2015) on 6 sessions for each participant, from which we used the first session only, as it could represent the whole dataset (the next sessions were repetition of the same stimuli to increase signal to noise ratio) and we preferred to avoid potential effect of extended familiarity with the stimuli on neural representations. The EEG data was collected during passive viewing (participants had no task but to keep fixating on the central fixation cross; Figure 1, bottom row) of 6 categories of objects with stimuli chosen from Kiani et al. (2007) as explained above. We used a pre-processed (i.e. band-pass-filtered in the range 0.03 to 50 Hz) version of the dataset which was available online^1^.

All the three datasets were collected at a sampling rate of 1000 Hz. For Datasets 1 and 2, only the trials which led to correct responses by participants, were used in the analyses. Each dataset consisted of data from 10 participants. Each object category in each dataset included 12 exemplars. To make the three datasets as consistent as possible, we pre-processed them differently from their original papers.

Specifically, the band-pass filtering range of Dataset 3 was 0.03 to 50 Hz, and we did not have access to the raw data to increase the upper cutting frequency to 200 Hz. Datasets 1 and 2 were band-pass-filtered in the range from 0.03 to 200 Hz before the data was split into trials. We also applied 50 Hz notch filters to Datasets 1 and 2 to remove line noise. Next, we generated different versions of the data by band-pass filtering the data in Delta (0.5-4 Hz), Theta (4-8 Hz), Alpha (8-12 Hz), Beta (12-16 Hz), Gamma (16-200Hz) bands to see if there is any advantage for the suggested Theta or Delta frequency bands (Watrous et al., 2015; Behroozi et al., 2016; Wang et al., 2018). We used finite-impulse-response (FIR) filters with 12 dB roll-off per octave for band-pass filtering of Datasets 1 and 2 and when evaluating the sub-bands of the three datasets. All the filters were applied before splitting the data into trials.

We did not remove artifacts (e.g. eye-related and movement-related) from the signals, as we and others have shown that sporadic artifacts have minimal effect in multivariate decoding (Grootswagers et al., 2017). To increase signal to noise ratios in the analyses, each unique stimulus had been presented to the participants 3, 6 and 12 times in Datasets 1, 2 and 3, respectively. Trials were defined in the time window from 200 ms before to 1000 ms after the stimulus onset to cover most of the range of event-related neural activations. The average pre-stimulus (−200 to 0 ms relative to the stimulus onset) signal amplitude was removed from each trial of the data. For more information about each dataset see Table 1 and the references to their original publications.

### Features

EEG signals are generated by inhibitory and excitatory post-synaptic potentials of cortical neurons. These potentials extend to the scalp surface and are recorded through electrodes as amplitudes of voltage in units of microvolts. Researchers have been using different aspects of these voltage recordings to obtain meaningful information about human brain processes. The main focus of this study is to compare the information content of features which are sensitive to temporal variabilities of neural activations against the “mean” of activity within the analysis window, which is conventionally used in decoding analysis (Grootswagers et al., 2017). Below we explain the mathematical formulas for each individual feature used in this study. We also provide brief information about potential underlying neural mechanisms which can lead to the information content provided by each feature.

We classified the features into five classes based on their mathematical similarity to simplify the presentation of the results and their interpretations. The five classes consist of Moment, Complexity, ERP, Frequency-domain and Multi-valued features. However, the classification of the features is not strict and the features might be classified based on other criteria and definitions. For example, complexity itself has different definitions (Tononi et al., 1998), such as degree of randomness, or degrees of freedom in a large system of interacting elements. There are also recent studies which split the variability features into the three categories of variance-, frequency- and information theory-based categories (Waschke et al., 2021). Therefore, each definition may exclude or include some of our features in the class. It is of note that, we only used the features which were previously used to decode categories of evoked potentials from EEG signals through multivariate decoding analysis. Nonetheless, there are definitely other features available, especially, those extracted from EEG time series collected during long-term monitoring of human neural representations in health and disorder (Fulcher and Jones, 2017). In presenting the features’ formulas, we avoided repeating the terms from the first feature to the last one. Therefore, the reader might need to go back a few steps/features to find the definitions of the terms. Note that, in this study, the analyses are performed in either 1000 ms time windows (i.e. number of samples used for feature extraction: *N*=1000) in the whole-trial analysis or 50 ms time windows (*N*=50) in time-resolved analysis.

### Moment features

These features are the most straightforward and intuitive features from which we might be able to extract information about neural processes. Mean, Variance, Skewness and Kurtosis are the 1^st^ to 4^th^ moments of EEG time series and can provide information about the shape of the signals and their deviation from stationarity, which is the case in evoked potentials (Rasoulzadeh et al., 2016; Wong et al., 2006). These moments have been shown to be able to differentiate visually evoked responses (Pouryzdian and Erfaninan, 2010; Alimardani et al., 2018). The 2^nd^ to 4^th^ moments are also categorized as variance-based features in recent studies (Waschke et al., 2021).

#### Mean

Mean amplitude of an EEG signal changes in proportion to the neural activation of the brain. It is by far the most common feature of the recorded neural activations used in analyzing brain states and cognitive processes both in univariate and multivariate analyses (Vidal et al., 2010; Hebart and Baker, 2017; Grootswagers et al., 2017; Karimi-Rouzbahani et al., 2019). In EEG, the brain activation is reflected as the amplitude of the recorded voltage across each electrode and the reference electrode at specific time points. To calculate the Mean feature, which is the first moment in statistics, the sample mean is calculated for each recorded EEG time series as:

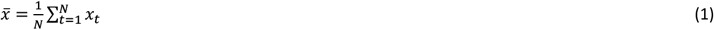

where 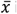 is the mean of the *N* time samples contained in the analysis window and *x*_*t*_refers to the amplitude of the recorded sample at time point *t. N*can be as small as unity as in the case of time-resolved EEG analysis (Grootswagers et al., 2017) or as large as it can cover the whole trial in whole-trial analysis. Accordingly, we set *N*=1000 (i.e. 1000 ms) and *N*= 50 (i.e. 50 ms) for the whole-trial and time-resolved decoding analyses, respectively.

#### Median

Compared to the Mean feature, Median is less susceptible to outliers (e.g. spikes) in the time series, which might not come from neural activations but rather from artifacts caused by the recording hardware, preprocessing, eye-blinks, etc. Median is calculated as:

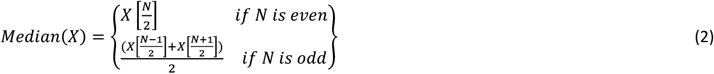

where *X* is the ordered values of samples in the time series *x*_*t*_for *t*=1,…,*N*.

#### Variance

Variance of an EEG signal is one simplest indicators showing how much the signal is deviated from stationarity i.e. deviated from its original baseline statistical properties (Wong et al., 2006). It is a measure of signal variabilities (within-trial here), has been shown to decline upon the stimulus onset potentially as a result of neural co-activation and has provided information about object categories in a recent EEG decoding study (Karimi-Rouzbahani et al., 2017a). Variance is calculated as:

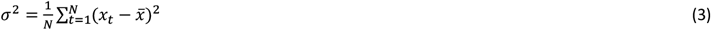

#### Skewness

While Variance is silent about the direction of the deviation from the mean, Skewness, which is the third signal moment, measures the degree of asymmetry in the signal’s probability distribution. In symmetric distribution (i.e. when samples are symmetric around the mean) skewness is zero. Positive and negative skewness indicates right- and left-ward tailed distribution, respectively. As the visually evoked ERP responses usually tend to be asymmetrically deviated in either positive or negative direction, even after baseline correction (Mazaheri and Jensen, 2008), we assume that Skewness should provide information about the visual stimulus if each category modulates the deviation of the samples differentially.

Skewness is calculated as:

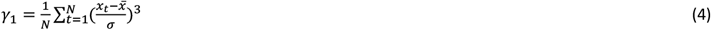

#### Kurtosis

Kurtosis reflects the degree of “tailedness” or “flattedness” of the signal’s probability distribution. Accordingly, the more heaviness in the tails, the less value of the Kurtosis and vice versa. Based on previous studies, Kurtosis has provided distinct representations corresponding to different classes of visually evoked potentials (Alimardani et al., 2018; Pouryzdian and Erfaninan, 2010). We test to see if Kurtosis plays a more generalized role in information coding e.g. coding of semantic aspects of visual information as well. It is the fourth standardized moment of the signal defined as:

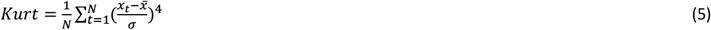

### Complexity features

There can potentially be many cases in which simple moment statistics such as Mean, Median, Variance, Skewness and Kurtosis, which rely on distributional assumptions, provide equal values for distinct time series (e.g. series A: 10, 20, 10, 20, 10, 20, 10, 20 vs. series B: 20, 20, 20, 10, 20, 10, 10, 10) for both of which the five above-mentioned features provide equal results. Therefore, we need more complex and possibly nonlinear measures which can detect subtle but meaningful temporal patterns from time series. The analysis of nonlinear signal features has recently been growing, following the findings showing that EEG reflects weak but significant nonlinear structures (Stam, 2005; Stepien, 2002).

Importantly, many studies have shown that the complexity of EEG time series can significantly alter during cognitive tasks such as visual (Bizas et al., 1999) and working memory tasks (Sammer et al., 1999; Stam, 2000). Therefore, it was necessary to evaluate the information content of nonlinear features for our decoding of object categories. As mentioned above, the grouping of these nonlinear features as “complexity” here is not strict and the features included in this class are those which capture complex and nonlinear patterns across time series. Although the accurate detection of complex and nonlinear patterns generally need more time samples compared to linear patterns (Procaccia, 1988), it has been shown that nonlinear structures can be detected from short EEG time series as well (i.e. through fractal dimensions; Preißl et al., 1997). Nonetheless, we extract these features from both time-resolved (50 samples) and whole-trial data (1000 samples) to ensure we do not miss potential information represented in longer temporal scales.

#### Lempel-Ziv complexity (LZ Cmplx)

Lempel-Ziv complexity measures the complexity of time series (Lempel et al., 1976). Basically, the algorithm counts the number of unique sub-sequences within a larger binary sequence. Accordingly, a sequence of samples with a certain regularity does not lead to a large LZ complexity. However, the complexity generally grows with the length of the sequence and its irregularity. In other words, it measures the generation rate of new patterns along a digital sequence. In a comparative work, it was shown that, compared to many other frequency metrics of time series (e.g. noise power, stochastic variability, etc.), LZ complexity has the unique feature of providing a scalar estimate of the bandwidth of time series and the harmonic variability in quasi-periodic signals (Aboy et al., 2006). It is widely used in biomedical signal processing and has provided successful results in the decoding of visual stimuli from neural responses in primary visual cortices (Szczepański et al., 2003). We used the code by Quang Thai^2^ implemented based on “exhaustive complexity” which is considered to provide the lower limit of the complexity as explained by Lempel et al. (1976). We used the signal median as a threshold to convert the signals into binary sequences for the calculation of LZ complexity. The LZ complexity provided a single value for each signal time series.

#### Fractal dimension

In signal processing, fractal is an indexing technique which provides statistical information about the complexity of time series. A higher fractal value indicates more complexity for a sequence as reflected in more nesting of repetitive sub-sequences at all scales. Fractal dimensions are widely used to measure two important attributes: self-similarity and the shape of irregularity. A growing set of studies have been using fractal analyses for the extraction of information about semantic object categories (such as living and non-living categories of visual objects; Ahmadi-Pajouh et al., 2018; Torabi et al., 2017) as well as simple checkerboard patterns (Namazi et al., 2018) from visually evoked potentials. In this study, we implemented two of the common methods for the calculation of fractal dimensions of EEG time series, which have been previously used to extract information about object categories as explained below. We used the implementations by Jesús Monge Álvarez^3^ for fractal analysis.

##### Higuchi’s fractal dimension (Higuchi FD)

In this method (Higuchi et al., 1988), a set of sub-sequences 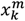 is generated in which *k* and *m* refer to the step size and initial value, respectively. Then, the length of this fractal dimension is calculated as:

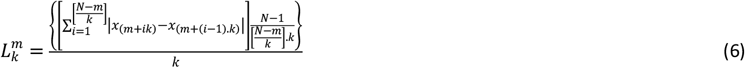

*where* 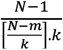 is the normalization factor. The length of the fractal curve at step size of *k* is calculated by averaging *k* sets of 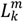. Finally, the resultant average will be proportional to *k*^−*D*^ where *D* is the fractal dimension. We set the free parameter of *k*equal to half the length of signal time series in the current study.

##### Katz’s fractal dimension (Katz FD)

We also calculated fractal dimension using the Katz’s method (Katz, 1988) as it showed a significant amount of information about object categories in a previous study (Torabi et al., 2017). The fractal dimension (*D*) is calculated as:

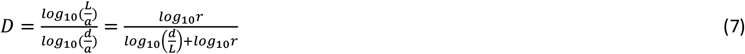

*where L* and *a* refer to the sum and average of the consecutive signal samples, respectively. Also *d* refers to the maximum distance between first sample and *i*^*th*^ sample of the signal which has the maximum distance from first sample as:

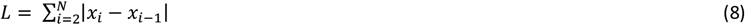

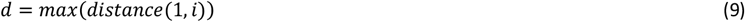

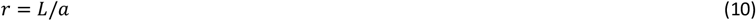

#### Hurst exponent (Hurst Exp)

Hurst exponent is widely used to measure the long-term memory in time-dependent random variables such as biological time series (Racine, 2011). In other words, it measures the degree of interdependence across samples in the time series and operates like an autocorrelation function over time. Hurst values between 0.5 and 1 suggest consecutive appearance of high signal values on large time scales while values between 0 and 0.5 suggest frequent switching between high and low signal values. Values around suggest no specific patterns among samples of a time series. It is defined as an asymptotic behavior of a rescaled range as a function of the time span of the time series defined as:

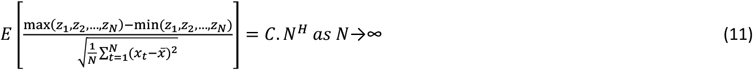

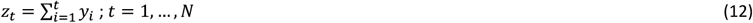

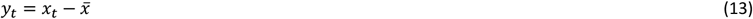

where *E* is the expected value, *C* is a constant and *H* is the Hurst exponent (Racine, 2011). We used the open-source implementation of the algorithm^4^, which has also been used previously for the decoding of object category information in EEG (Torabi et al., 2017).

#### Entropy

Entropy can measure the perturbation in time series (Waschke et al., 2021). A higher value for entropy suggests a higher irregularity in the given time series. Precise calculation of entropy usually requires considerable number of samples and is also sensitive to noise. Here we used two methods for the calculation of entropy, each of which has its advantages over the other.

##### Approximate entropy (Apprx Ent)

Approximate entropy was initially developed to be used for medical data analysis (Pincus and Huang, 1992), such as heart rate, and then was extended to other areas such as brain data analysis. It has the advantage of requiring a low computational power which makes it perfect for real-time applications on low sample sizes (<50). However, the quality of this entropy method is impaired on lower lengths of the data. This metric detects changes in episodic behavior which are not represented by peak occurrences or amplitudes (Pincus and Huang, 1992). We used an open-source code^5^ for calculating approximate entropy. We set the embedded dimension and the tolerance parameters to 2 and 20% of the standard deviation of the data respectively, to roughly follow a previous study (Shourie et al., 2014) which compared approximate entropy in visually evoked potentials and found differential effects across artist vs. non-artist participants when looking at paintings.

##### Sample entropy (Sample Ent)

Sample entropy, which is a refinement of the approximate entropy, is frequently used to calculate regularity of biological signals (Richman et al., 2000). Basically, it is the negative natural logarithm of the conditional probability that two sequences (subset of samples), which are similar for *m* points remain similar at the next point. A lower sample entropy also reflects a higher self-similarity in the time series. It has two main advantages to the approximate entropy: it is less sensitive to the length of the data and is simpler to implement. However, it does not focus on self-similar patterns in the data. We used the Matlab “entropy” function for the extraction of this feature, which has already provided category information in a previous study (Torabi et al., 2017). See (Richman et al., 2000; Subha et al., 2010) for the details of the algorithm.

#### Autocorrelation (Autocorr)

Autocorrelation determines the degree of similarity between the samples of a given time series and a time-lagged version of the same series. It detect periodic patterns in signals, which is an integral part of EEG time series. Therefore, following recent successful attempts in decoding neural information using the autocorrelation function from EEG signals (Wairagkar et al., 2016), we evaluated the information content of the autocorrelation function in decoding visual object categories. As neural activations reflect many repetitive patterns across time, the autocorrelation function can quantify the information contents of those repetitive patterns. Autocorrelation is calculated as:

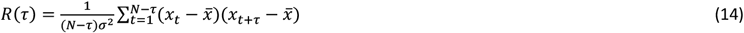

where *τ* indicates the number of lags in samples of the shifted signal. A positive value for autocorrelation indicates a strong relationship between the original time series and its shifted version, whereas a negative autocorrelation refers to an opposite pattern between them. Zero autocorrelation indicates no relationship between the original time series and its shifted version. In this study, we extracted autocorrelations for 30 consecutive lags ([*τ*=1, 2, …, 30]) used their average in classification. Please note that each lag refers to 1 ms as the data was sampled at 1000 Hz.

#### Hjorth parameters

Hjorth parameters are descriptors of statistical properties of signals introduced by Hjorth (1970). These parameters are widely used in EEG signal analysis for feature extraction across a wide set of applications including visual recognition (Joshi et al., 2018; Torabi et al., 2017). These features consist of Activity, Mobility and Complexity as defined below. As the Activity parameter is equivalent to the signal Variance, which we already explained, we do not repeat it.

##### Hjorth complexity (Hjorth Cmp)

It determines the variation in time series’ frequency by quantifying the similarity between the signal and a pure sine wave leading to a value of 1 in case of perfect match. In other words, values around 1 suggest lower complexity for a signal. It is calculated as:

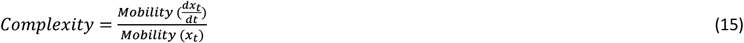

##### Hjorth mobility (Hjorth Mob)

It determines the proportion of standard deviation of the power spectrum as is calculated below, where *var* refers to the signal variance.

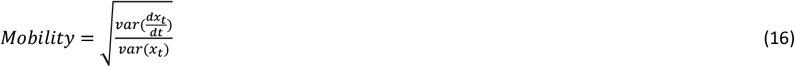

where *var* refers to the variance.

### ERP components (N1, P1, P2a and P2b)

An ERP is a measured brain response to a specific cognitive, sensory or motor event that provides an approach to studying the correlation between the event and neural processing. According to the latency and amplitude, ERP is split into specific sub-windows called components. Here, we extracted ERP components by calculating mean of signals in specific time windows to obtain the P1 (80 to 120 ms), N1 (120 to 200 ms), P2a (150 to 220 ms) and P2b (200 to 275 ms) components, which were shown previously to provide significant amounts of information about visual object and face processing in univariate (Rossion et al., 2000; Rousselett et al., 2007) and multivariate analyses (Chan et al., 2011; Jadidi et al., 2016; Wang et al., 2012). As these components are calculated in limited and specific time windows, in the whole-trial analysis, they reflect “Mean” of activity in their specific time windows, rather than the whole post-stimulus window. They will be also absent from time-resolved analyses by definition.

### Frequency-domain features

Neural variability is commonly analyzed in frequency domain by calculating spectral power across frequency bands. Specifically, as data transformation from time to frequency domain is almost lossless using Fourier transform, oscillatory power basically reflects frequency-specific variance (with the total power reflecting the overall variance of the time series (Waschke et al., 2021)). Motivated by previous studies showing signatures of object categories in the frequency domain (Behroozi et al., 2016; Rupp et al., 2017; Iranmanesh and Rodriguez-Villegas, 2017; Joshi et al., 2018; Jadidi et al., 2016) and the representation of temporal codes of visual information in the frequency domain (Eckhorn et al., 1988), we also extracted frequency-domain features to see if they could provide additional category-related information to time-domain features. It is of note that, while the whole-trial analysis allows us to compare our results with previous studies, the evoked EEG potentials are generally nonstationary (i.e. their statistical properties change along the trial), and potentially dominated by low-frequency components. Therefore, the use of time-resolved analysis, which looks at more stationary sub-windows of the signal (e.g. 50 samples here), will allow us to detect subtle high-frequency patterns of neural codes.

#### Signal power (Signal Pw)

Power spectrum density (PSD) represents the intensity or the distribution of the signal power into its constituent frequency components. This feature was motivated by previous studies showing associations between aspects of visual perception and power in certain frequency bands (Rupp et al., 2017; Behroozi et al., 2016; Majima et al., 2014). According to the Fourier analysis, signals can be broken into its constituent frequency components or a spectrum of frequencies in a specific frequency range. Here, we calculated signal power using the PSD as in (17).

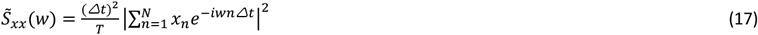

where *x*_*n*_ = *x*_*nΔt*_ is signal sampled at a rate of 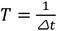 and *w* is the frequency at which the signal power is calculated. As signal power is a relatively broad term, including the whole power spectrum of the signal, we also extracted a few more parameters from the signal frequency representation to see what specific features in the frequency domain (if any) can provide information about object categories.

#### Mean frequency (Mean Freq)

Motivated by the successful application of mean and median frequencies in the analysis of EEG signals and their relationship to signal components in the time domain (Intrilligator and Polich, 1995; Abootalebi et al., 2009), we extracted these two features from the signal power spectrum to obtain a more detailed insight into the neural dynamics of category representations. Mean frequency is the average of the frequency components available in a signal. Assume a signal consisting of two frequency components of *f*_1_ and *f*_2_. The Mean frequency of this signal is 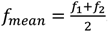. Generally, the mean normalized (by the intensity) frequency is calculated using the following formula:

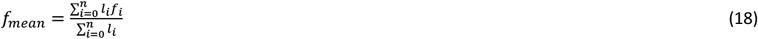

where *n* is the number of splits of the PSD, *f*_*i*_ and *l*_*i*_ are the frequency and the intensity of the PSD in its *i*^*th*^ slot, respectively. It was calculated using Matlab “meanfreq” function.

#### Median frequency (Med Freq)

Median frequency is the median normalized frequency of the power spectrum of a time-domain signal. It is calculated similarly to the signal median in the time domain, however, here the values are the power intensity in different frequency bins of the PSD. This feature was calculated using Matlab “medfreq” function.

#### Power and Phase at median frequency (Pw MdFrq and Phs MdFrq)

Interestingly, apart from the median frequency itself, which reflects the frequency aspect of the power spectrum, the power and phase of the signal at the median frequency have also been shown to be informative about aspects of human perception (Joshi et al., 2018; Jadidi et al., 2016). Therefore, we also calculated the power and phase of the frequency-domain signals at the median frequency as features.

#### Average frequency (Avg Freq)

Evoked potentials show a few number of positive and negative peaks after the stimulus onset, and they might show deviation in the positive or negative directions depending on the information content (Mazaheri and Jensen, 2008). Therefore, we also evaluated the Average (zero-crossing) frequency of the ERPs by counting the number of times the signal swapped signs during the trial. Note that each trial is baselined according to the average amplitude of the same trial in the last 200 ms immediately before the stimulus onset. We calculated the average frequency on the post-stimulus time window.

#### Spectral edge frequency (SEF 95%)

SEF is a common feature used in monitoring the depth of anesthesia and stages of sleep using EEG (Iranmanesh and Rodriguez-Villegas, 2017). It measures the frequency which covers X percent of the PSD. X is usually set between 75% to 95%. Here we set X to 95%. Therefore, this reflects the frequency observed in a signal which covers 95% of a signal power spectrum.

### Multi-valued features

The main hypothesis of the present study is that, we can potentially obtain more information about object categories as well as behavior if we take into account the temporal variability of neural activity within the analysis window (i.e. trial) rather than averaging the samples as in conventional decoding analyses. While the above variability-sensitive features return a single value from each individual time series (analysis window), a more flexible feature would allow as many informative patterns to be detected from an individual time series. Therefore, we extracted other features, which provide more than one value per analysis window, so that we can select the most informative values from across electrodes and time points simultaneously (see *Dimensionality reduction* below). We also included the Original Magnitude Data as our reference feature, so that we know how much (if at all) our feature extraction and selection procedures improved decoding.

#### Inter-electrode correlation (Cross Corr)

Following up on recent studies, which have successfully used inter-area correlation in decoding object category information from EEG activations (Majima et al., 2014; Karimi-Rouzbahani et al., 2017a; Tafreshi et al., 2019), we extracted inter-electrode correlation to measure the similarity between pairs of signals, here, from different pairs of electrodes. This feature of correlated variability, quantifies co-variability of neural activations across pairs of electrodes. Although closer electrodes tend to provide more similar (and therefore correlated) activation, compared to further electrodes (Hacker et al., 2017), the inter-electrode correlation can detect correlations which are functionally relevant and are not explained by the distance (Karimi-Rouzbahani et al., 2017a). This feature detects similarities in temporal patterns of fluctuations across time between pairs of signals, which. It is calculated as:

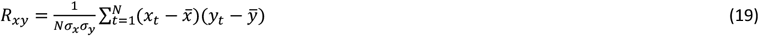

where *x* and *y* refer to the signals obtained from electrodes *x* and *y*, respectively. We calculated the cross-correlation between each electrode and all the other electrodes to form a cross-correlation matrix. Accordingly, we initially obtained all the unique possible pairwise inter-electrode correlations (465, 465 and 8128 unique values for Datasets 1, 2 and 3, respectively), which were then reduced in dimension using PCA to the equal number of dimensions obtained for single-valued features.

#### Wavelet transform (Wavelet)

Recent studies have shown remarkable success in decoding of object categories using the Wavelet transformation of the EEG time series (Taghizadeh-Sarabi et al., 2015; Torabi et al., 2017). Considering the time- and frequency-dependent nature of ERPs, Wavelet transform seems to be a very reasonable choice as it provides a time-frequency representation of signal components. It determines the primary frequency components and their temporal position in time series. The transformation passes the signal time series through digital filters (Guo et al., 2009; equation (20)), using the convolution operator, each of which adjusted to extract a specific frequency (scale) at a specific time as in (20):

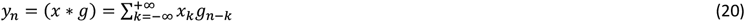

where *g* is the digital filter and * is the convolution operator. This filtering procedure is repeated for several rounds (levels) filtering low-(approximations) and high-frequency (details) components of the signal to provide more fine-grained information about the constituent components of the signal. This can lead to coefficients which can potentially discriminate signals evoked by different conditions. Following up on a previous study (Taghizadeh-Sarabi et al., 2015), and to make the number of Wavelet features comparable in number to signal samples, we used detail coefficients at five levels *D*1, …, *D*5 as well as the approximate coefficients at level 5, *A*5. This led to 1015 and 57 features in the whole-trial and in the 50 ms sliding time windows, respectively. We used the “*Symlet2”* basis function for our Wavelet transformations as implemented in Matlab.

#### Hilbert transform (Hilb Amp and Hilb Phs)

Hilbert transform provides amplitude and phase information about the signal and has recently shown successful results in decoding visual letter information from ERPs (Wang et al., 2018). The phase component of the Hilbert transform can qualitatively provide the spatial information obtained from the Wavelet transform leading to their similarity evaluating neuronal synchrony (Le Van Quyen et al., 2001). However, it is still unclear which method can detect category-relevant information from the nonstationary ERP components more effectively. Hilbert transform is described as a mapping function that receives a real signal *x*_*t*_(as defined above), and upon convolution with the function 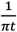, produces another function of a real variable *H*(*x*)(*t*) as:

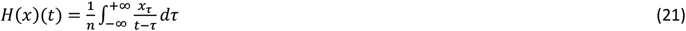

where *H*(*x*)(*t*) is a frequency-domain representation of the signal *x*_*t*_, which has simply shifted all the components of the input signal by 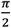. Accordingly, it produces one amplitude and one phase component per samples in the time series. In the current study, Hilbert transform was applied on 1000 and 50 samples in the whole-trial and time-resolved analysis, respectively. We used the amplitude and phase components separately to discriminate object categories in the analyses.

#### Amplitude- and Phase-locking (Amp Lock and Phs Lock)

Although inter-electrode correlated variability (*Cross Corr*), which is interpreted as inter-area connectivity, have successfully provided object category information (Majima et al., 2014; Karimi-Rouzbahani et al., 2017a), previous studies suggested that neural communication is realized through amplitude- and phase-locking/coupling (Bruns et al., 2000; Siegel et al., 2012; Engel et al., 2013). More recently, researchers have quantitatively shown that amplitude- and phase-locking detect distinct signatures of neural communication across time and space from neural activity (Siems and Siegel, 2020; Mostame and Sadaghiani, 2020). Therefore, in line with recent studies, which successfully decoded object categories using inter-area correlated variability as neural codes (Tafreshi et al., 2019), we extracted amplitude- and phase-locking as two major connectivity features which might contain object category information as well. Briefly, amplitude-locking refers to the coupling between the envelopes of two signals (electrodes) and reflects the correlation of activation amplitude. To estimate the amplitude locking between two signals, we extracted the envelopes of the two signals using Hilbert transform (Gabor, 1946; explained below), then estimated the Pearson correlation between the two resulting envelopes as amplitude locking.

Phase locking, on the other hand, refers to the coupling between the phases of two signals and measures the synchronization of rhythmic oscillation cycles. To measure phase locking we used one of the simplest implementations, the phase locking value (PLV), which includes minimal mathematical assumptions (Bastos and Schoffellen, 2016) calculated as below:

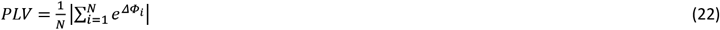

where *N* is the number of trials and *ΔΦ* is the phase difference between the signals to electrode pairs. As we used multivariate decoding without any trial-averaging, *N* was equal to 1 here. The calculation of amplitude and phase locking was performed on all electrode pairs leading to 465 and 8128 unique numbers for the 31- (Datasets 1 and 2) and 128-electrode (Dataset 3) datasets before dimension reduction was performed.

#### Original magnitude data (Orig Mag)

We also used the post-stimulus original magnitude data (i.e. 1000 or 50 samples for the whole-trial and sliding time windows, respectively) to decode object category information without any feature extraction. This provided a reference to compare the information content of the Mean and variability features to see if the former provided any extra information.

### Multivariate decoding

We used multivariate decoding to extract information about object categories from our EEG datasets. Basically, multivariate decoding, which has been dominating neuroimaging studies recently (Haynes et al., 2015; Grootswagers et al., 2017; Hebart and Baker, 2018), measures the cross-condition dissimilarity/contrast to quantify information content in neural representations. We used linear discriminant analysis (LDA) classifiers in multivariate analysis to measure the information content across all possible pairs of object categories within each dataset. Specifically, we trained and tested the classifiers on e.g. animal vs. car, animal vs. face, animal vs. plane, car vs. plane, face vs. car and plane vs. face categories, then averaged the 6 decoding results and reported them for each participant. The LDA classifier has been shown to be robust when decoding object categories from M/EEG (Grootswagers et al., 2017; Grootswagers et al., 2019), has provided higher decoding accuracies than Euclidean distance and Correlation based decoding methods (Carlson et al., 2013) and was around 30 times faster to train in our initial analyses compared to the more complex classifier of Support-Vector Machine (SVM). We ran our initial analysis and found similar results for the LDA and SVM, and used LDA to save the time. We used a 10-fold cross-validation procedure in which we trained the classifier on 90% of the data and tested it on the left-out 10% of the data, repeating the procedure 10 times until all trials from the pair of categories participate once in the training and once in the testing of the classifiers. We repeated the decoding across all possible pairs of categories within each dataset, which were 6, 6 and 15 pairs for Datasets 1, 2 and 3, which consisted of 4, 4 and 6 object categories, respectively. Finally, we averaged the results across all combinations and reported them as the average decoding for each participant.

In the whole-trial analyses, we extracted the above-mentioned features from the 1000 data samples after the stimulus onset (i.e. from 1 to 1000 ms). In the time-resolved analyses, on the other hand, we extracted the features from 50 ms sliding time windows in steps of 5 ms across the time course of the trial (−200 to 1000 ms relative to the stimulus onset time). Therefore, in time-resolved analyses, the decoding rates at each time point reflect the results for the 50 ms window around the time point, from - 25 to +24 ms relative to the time point. Time-resolved analyses allowed us to evaluate the evolution of object category information across time as captured by different features.

### Dimensionality reduction

The multi-valued features (e.g. inter-electrode correlation, wavelet, Hilbert amplitude and phase, Amplitude and Phase locking and Original magnitude data) resulted in more than a single feature value per trial per sliding time window. This could provide higher decoding values compared to the decoding values obtained from single-valued features merely because of including a higher number of features. Moreover, when the features outnumber the observations (i.e. trials here), the classification algorithm can over-fit to the data (Duda et al., 2012). Therefore, to obtain comparable decoding accuracies across single-valued and multi-valued features and to avoid potential over-fitting of classifier to the data we used principle component analysis (PCA) to reduce the dimension of the data in multi-valued features. Accordingly, we reduced the number of the values in the multi-valued features to ***one*** per time window per trial, which equaled the number of values for the single-valued features. To avoid potential leakage of information from testing to training (Pulini et al., 2019), we applied the PCA algorithm on the training data (folds) only and used the training PCA parameters (i.e. eigen vectors and means) for both training and testing sets for dimension reduction in each cross-validation run separately. We only applied the dimension-reduction procedure on the multi-valued features. Note that, we did not reduce the dimension of the neural space (columns in the dimension-reduced data matrix) to below the number of electrodes “*e*” (opposite to Hatamimajoumerd et al., 2019) as we were interested in qualitatively comparing our results with the vast literature currently using multivariate decoding with all sensors (Grootswagers et al., 2017; Karimi-Rouzbahani et al., 2018; Hebart and Baker 2017). Also, we did not aim at finding more than one feature per trial, per time window, as we wanted to compare the results of multi-valued features with those of single-valued features, which only had a single value per trial, per time window.

One critical point here is that, we applied the PCA on ***the concatenated data from all electrodes and values obtained from each individual feature*** (e.g. wavelet coefficients in Wavelet), within each analysis window (e.g. 50 ms in time-resolved decoding). Therefore, for the multi-valued features, the “e” selected dimensions were the most informative ***spatial*** and ***temporal*** patterns detected across both *electrodes* and *time samples*. Therefore, it could be the case that, within a given time window, two of the selected dimensions were from the same electrode (i.e. because two elements from the same electrode were more informative than the other electrode), which would lead to some electrodes not having any representatives among the selected dimensions. This is in contrast to the single-valued features (e.g. Mean) from which we only obtained one value per analysis window per electrode, limiting the features to only the ***spatial*** patterns within the analysis window, rather than both spatial and temporal patterns.

### Statistical analyses

#### Bayes factor analysis

As in our previous studies (Grootswagers et al., 2019; Robinson et al., 2019), to determine the evidence for the null and the alternative hypotheses, we used Bayes analyses as implemented by Bart Krekelberg based on Rouder et al. (2012). We used standard rules of thumb for interpreting levels of evidence (Lee and Wagenmakers, 2014; Dienes, 2014): Bayes factors of >10 and <1/10 were interpreted as strong evidence for the alternative and null hypotheses, respectively, and >3 and <1/3 were interpreted as moderate evidence for the alternative and null hypotheses, respectively. We considered the Bayes factors which fell between 3 and 1/3 as suggesting insufficient evidence either way.

In the whole-trial decoding analyses, we asked whether there was a difference between the decoding values obtained from all possible pairs of features and also across frequency bands within every feature. Accordingly, we performed the Bayes factor analysis and calculated the Bayes factors as the probability of the data under alternative (i.e. difference) relative to the null (i.e. no difference) hypothesis between all possible pairs of features and also across frequency bands within every feature and dataset separately. The same procedure was used to evaluate evidence for difference (i.e. alternative hypothesis) or no difference (i.e. null hypothesis) in the maximum and average decoding accuracies, the time of maximum and above-chance decoding accuracies across features for each dataset separately.

We also evaluated evidence for the alternative of above-chance decoding accuracy vs. the null hypothesis of no difference from chance. For that purpose, we performed Bayes factor analysis between the distribution of actual accuracies obtained and a set of 1000 accuracies obtained from random permutation of class labels across the same pair of conditions (null distribution) on every time point (or only once for the whole-trial analysis), for each feature and dataset separately. No correction for multiple comparisons was performed when using Bayes factors as they are much more conservative than frequentist analysis in providing false claims with confidence (Gelman and Tuerlinckx, 2000; Gelman et al., 2012). The reason for the less susceptibility of Bayesian analysis compared to classical statistics, is the use of priors, which if chosen properly (here using the data-driven approach developed by Rouder et al. (2012)), significantly reduce the chance of making type I (false positive) errors.

The priors for all Bayes factor analyses were determined based on Jeffrey-Zellner-Siow priors (Jeffreys, 1961; Zellner and Siow, 1980) which are from the Cauchy distribution based on the effect size that is initially calculated in the algorithm (Rouder et al., 2012). The priors are data-driven and have been shown to be invariant with respect to linear transformations of measurement units (Rouder et al., 2012), which reduces the chance of being biased towards the null or alternative hypotheses.

#### Random permutation testing

To evaluate the significance of correlations between decoding accuracies and behavioral reaction times, we calculated the percentage of the actual correlations that were higher (when positive) or lower (when negative) than a set of 1000 randomly generated correlations. These random correlations were generated by randomizing the order of participants’ data in the behavioral reaction time vector (null distribution) for every time point and feature separately. The true correlation was considered significant if it surpassed 95% of the randomly generated correlations in the null distribution in either positive or negative directions (p < 0.05) and the p-values were corrected for multiple comparisons across time using Matlab mafdr function which works based on fix rejection region (Storey, 2002).

## Results

To check the information content of different features of the EEG activity about object categories, we performed multivariate pattern decoding on both the whole-trial as well as time-resolved data. The whole-trial analysis was aimed at providing results comparable to previous studies most of which performed whole-trial analysis. The time-resolved analysis, however, was the main focus of the present study and allowed us to check the information and temporal dynamics of variability-based neural codes as captured by different features. In figures 2 and 3, we only present a summary of the results with emphasis on the comparison between the time-specific ERP components, the most informative features detecting neural variability (i.e. Wavelet and Orig Mag), and the conventional Mean feature, which ignores potential information in neural variabilities. The complete comparison between the 32 features are provided in Supplementary materials, but briefly explained in the manuscript.

**Figure 2.**
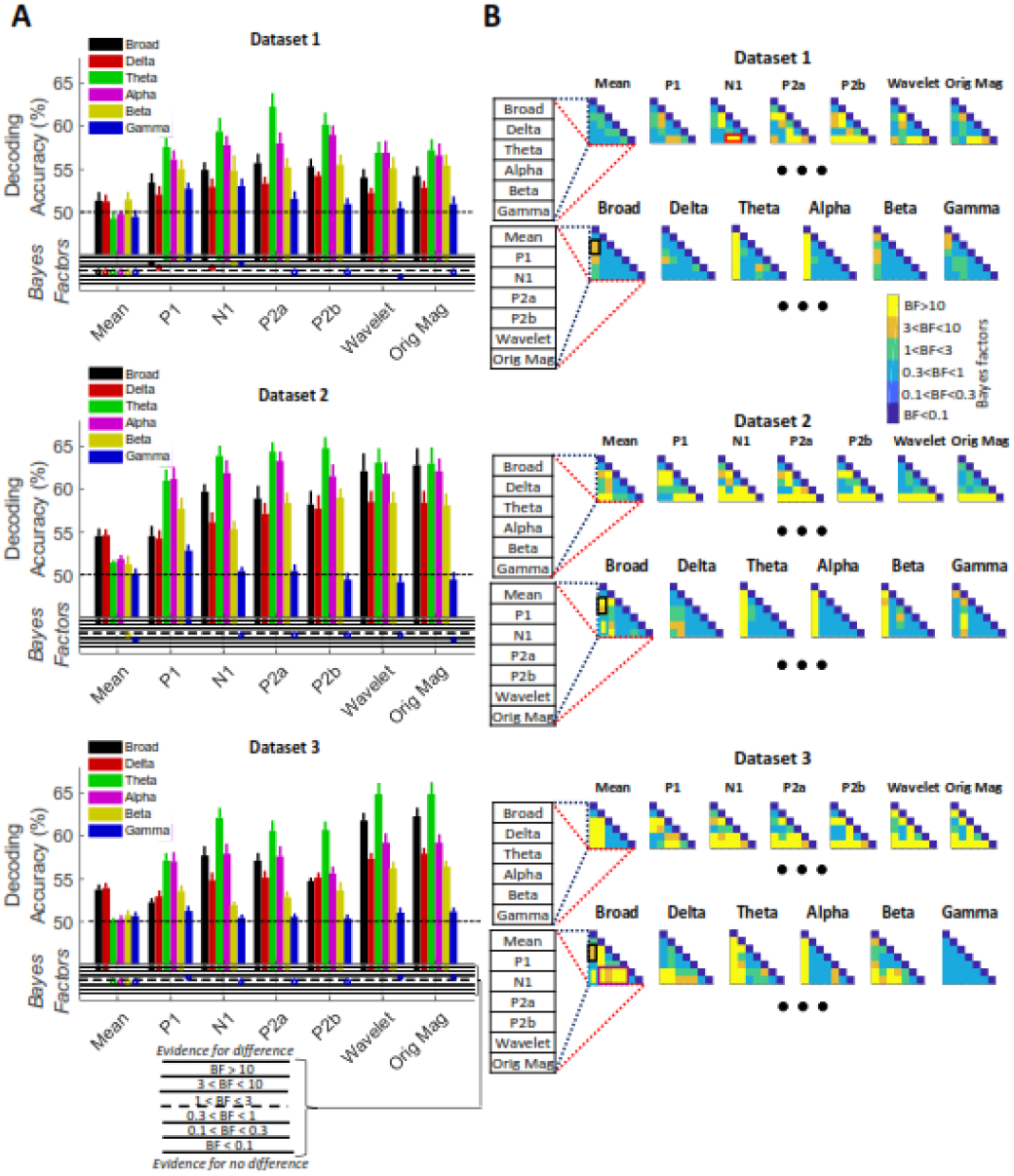
Whole-trial decoding of object categories in the three datasets across the Broad-band and different frequency bands (A) with their Bayesian analyses (B). The results are only presented for features of Mean, ERP components, Wavelet and Orig Mag. For full results including other features see Supplementary Figures 1 and (A) The black horizontal dashed lines on the top panels refer to chance-level decoding. Thick bars show the average decoding across participants (error bars Standard Error across participants). Bayes Factors are shown in the bottom panel of each graph: Filled circles show moderate/strong evidence for either hypothesis and empty circles indicate insufficient evidence. They show the results of Bayes factor analysis when evaluating the difference from chance-level decoding. (B) Top panel Bayes matrices compare the decoding results within each frequency band, across features separated by datasets. Bottom panel Bayes matrices compare decoding results across different frequency bands and dataset separately. Colors indicate different levels of evidence for existing difference (moderate 3<BF<10, Orange; strong BF>10, Yellow), no difference (moderate 0.1<BF<0.3, light blue; strong BF<0.1, dark blue) or insufficient evidence (1<BF<3 green; 0.3<BF<1 Cyan) for either hypotheses. For example, for Dataset 1, there is strong evidence for higher decoding values for the N1 feature in the Theta and Alpha band than in Gamma band as indicated by the red box.

**Figure 3.**
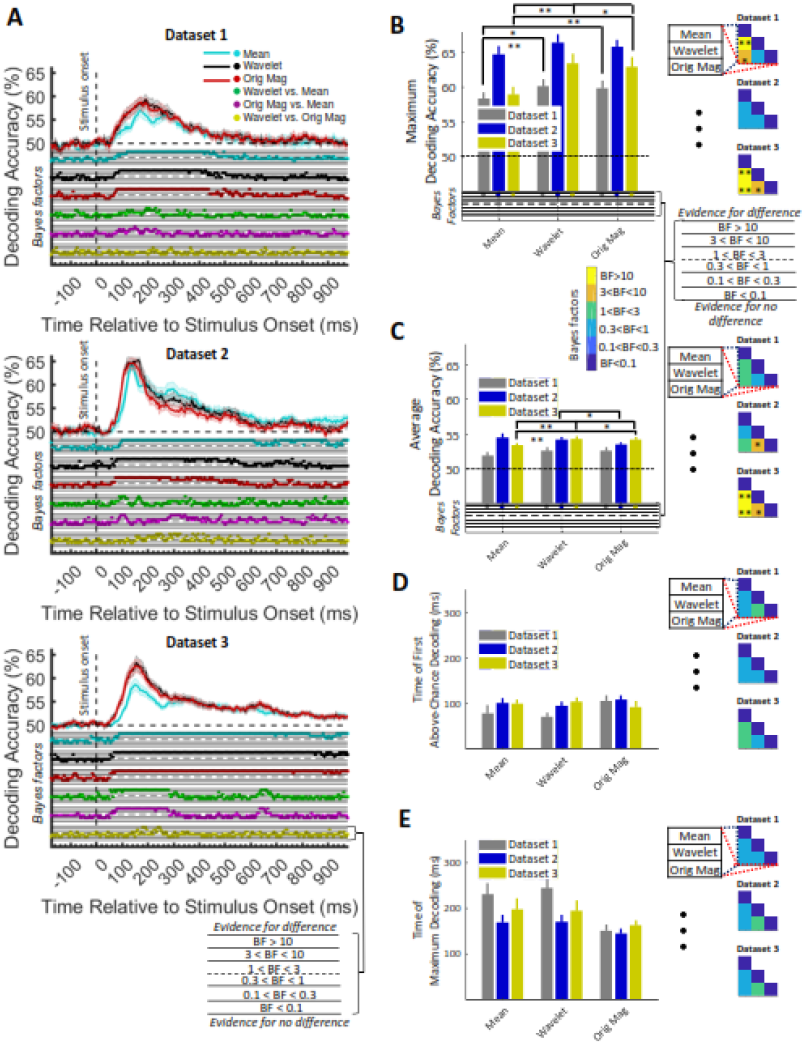
Time-resolved decoding of object categories from the three datasets for 3 of the target features (A) and their extracted timing and amplitude parameters (B-E). (A) Top section in each panel shows the decoding accuracies across time and the bottom section shows the Bayes factor evidence for the difference of the decoding accuracy compared to chance-level decoding. The solid lines show the average decoding across participants and the shaded area the Standard Error across participants. The horizontal dashed lines on the top panel refer to chance-level decoding. Filled circles in the Bayes Factors show moderate/strong evidence for either difference or no difference from chance-level or across features and empty circles indicate insufficient evidence for either hypotheses. (B) Timing and amplitude parameters extracted from the time-resolved accuracies in (A). (B-E) Left: the maximum and average decoding accuracies, the time of maximum and the first above-chance decoding. The horizontal dashed lines refer to chance-level decoding. Thick bars show the average across participants (error bars Standard Error across participants). Bottom section on (B) and (C) show the Bayes factor evidence for the difference of the decoding accuracy compared to chance-level decoding. (B-E) Right: matrices compare the parameters obtained from different features. Different levels of evidence for existing difference (moderate 3<BF<10, Orange; strong BF>10, Yellow), no difference (moderate 0.1<BF<0.3, light blue; strong BF<0.1, dark blue) or insufficient evidence (1<BF<3 green; 0.3<BF<1 Cyan) for either hypotheses. Filled circles in the Bayes Factors show moderate/strong evidence for either hypothesis and empty circles indicate insufficient evidence. Single and double stars indicate moderate and strong evidence for difference between the parameters obtained from decoding curves of the three features.

### Can the features sensitive to temporal variabilities, provide additional category information to the conventional “mean” feature

To answer the first question, we compared decoding accuracies in the whole-trial time span (0 to 1000 ms relative to stimulus onset) across all features and for each dataset separately (see the complete results in Supplementary Figures 1 and 2 and summary results in Figure 2, black bars). There was not enough (BF>3) evidence for above-chance decoding for majority of features (e.g. moment features, complexity and frequency-domain features, Supplementary Figure 1; black bars and their Bayesian analyses). However, consistently across the three datasets, there was moderate (3<BF<10) or strong (BF>10) evidence for above-chance decoding for all ERP components (N1, P1, P2a and P2b), Wavelet coefficients (Wavelet) and Original magnitude data (Orig Mag), which were either targeted at specific time windows within the trial (i.e. ERPs) or could detect temporal variabilities within the trial (i.e. Wavelet and Orig Mag; Figure 2A; black bars).

Importantly, in all three datasets, there was moderate (3<BF<10) or strong (BF>10) evidence that ERP components of N1 and P2a provided *higher* decoding values than the Mean (Figure 2B; black boxes in Bayes matrices). There was also strong evidence (BF>10), that the Wavelet and Orig Mag features outperformed the Mean feature in datasets 2 and 3 (Figure 2B; blue boxes in Bayes matrices). This shows that simply using the earlier ERP components of N1 and P2a can provide more information than using the Mean activity across the whole trial. This was predictable, as the Mean across the whole trial simply ignores within-trial temporally specific information. Interestingly, even ERPs were outperformed by Wavelet and Orig Mag features in Dataset 3 (but not the opposite across the 3 datasets; Figure 2B; violet boxes in Bayes matrices). This suggests that, even further targeting the most informative elements (i.e. Wavelet), and/or data samples (i.e. Orig Mag) within the trial can lead to improved decoding. Note that, the Wavelet and Orig Mag features provided the most informative temporal patterns/samples upon the dimension reduction procedure applied on their extracted features (see *Methods*).

Following previous observations about the advantage of Delta (Watrous et al., 2015; Behroozi et al., 2016) and Theta (Wang et al., 2018) frequency bands, we compared the information content in the Delta (0.5-4 Hz), Theta (4-8 Hz), Alpha (8-12 Hz), Beta (12-16 Hz), Gamma (16-200Hz) and Broad frequency bands. We predicted the domination of Theta frequency band, following suggestions about the domination of Theta frequency band in feed-forward visual processing (Bastos et al., 2015). For our top-performing ERP, Wavelet and Orig Mag features, we saw consistent domination of Theta followed by the Alpha frequency band (Figure 2A). Interestingly, for the ERP components, the decoding in Theta band even outperformed the Broad band (BF>3 for P2b), which contained the whole frequency spectrum. Note that, as opposed to previous suggestions (Karakaş et al., 2000), the domination of the Theta frequency band in ERP components could not be trivially predicted by their timing relative to the stimulus onset. If this was the case here, the P2b component (200 to 275 ms) should have elicited its maximum information in the Delta (0.5 to 4Hz) and Theta (4-8 Hz), rather than the Theta and Alpha (8-12 Hz) frequency bands. For the Mean feature, on the other hand, the Delta band provided the highest information level, comparable to the level of the Broad-band activity. This confirms that Broad-band whole-trial Mean activity, reflects the general trend of the signal (low-frequency component).

Together, we observed that the features which are targeted at informative windows of the trial (ERP components), and those sensitive to informative temporal variabilities (Wavelet and Orig Mag) could provide additional category information to the conventionally used Mean of activity. We observed that Theta frequency band, which has been suggested to support feed-forward information flow, is also dominant in our datasets, which are potentially dominated by feed-forward processing of visual information during object perception. Next, we will compare the temporal dynamics of information encoding across our features.

### Do the features sensitive to temporal variabilities evolve over similar time windows to the “mean” feature

One main insight that EEG decoding can provide is to reveal the temporal dynamics of cognitive processes. However, the Mean activity, which has dominated the literature (Grootswagers et al., 2017), might hide or distort the true temporal dynamics as it ignores potentially informative temporal variabilities (codes) within the analysis window. Therefore, we systematically compared the information content of a large set of features which are sensitive to temporal variabilities using time-resolved decoding (50 ms sliding time windows in steps of 5 ms; see the rationale for choosing the 50 ms windows in Supplementary Figure 3A). By definition, we do not have the time-resolved decoding results for the ERP components here.

Before presenting the time-resolved decoding results, to validate the results and suggestions made about our whole-trial decoding (Figure 2), we performed two complementary analyses. First, we checked to see if the advantage of the Theta-to Broad-band decoding in the whole-trial analysis (Figure 2), could generalize to time-resolved decoding: we observed the same effect in the (variability-sensitive) Wavelet feature (in many time points especially for Dataset 2; BF>3), but not in the (variability-insensitive) Mean feature (Supplementary Figure 3B). This could possibly be explained by the smoothing (low-pass filtering) effect of the Mean feature making both Theta- and Broad-band data look like low-frequency data. Next, we utilized the spatiotemporal specificity of classifier weights and time-resolved decoding to see if Theta-band information would show a feed-forward trend on the scalp to support our earlier suggestion. Visual inspection suggests information spread from posterior to anterior parts of the scalp (Supplementary Figure 4), supporting the role of Theta-band activity in feed-forward processing. Despite these observations, we used Broad-band signals in the following analyses to be able to compare our results with previous studies, which generally used the Broad-band activity.

Time-resolved decoding analyses showed that for all features, including the complexity features, which were suggested to need large sample sizes (Procaccia, 2000), there was moderate (3<BF<10) or strong (BF>10) evidence for above-chance decoding at some time points consistently across the three datasets (Supplementary Figures 5A). However, all features showed distinct temporal dynamics to each other and across datasets. The between-dataset dissimilarities, could be driven by many dataset-specific factors, including duration of image presentation (Carlson et al., 2013). However, there were also similarities between the temporal dynamics of different features. For example, the time points of first strong (BF>10) evidence for above-chance decoding ranged from 75 ms to 195 ms (Supplementary Figure 5A and E) and the decoding values reached their maxima in the range between 150 ms to 220 ms (Supplementary Figures 5A and D) across features. This is consistent with many decoding studies showing the temporal dynamics of visual processing in the brain (Isik et al., 2013; Cichy et al., 2014; Karimi-Rouzbahani et al., 2021b). There was no feature which consistently preceded or followed other features, to suggest the existence of very early or late neural codes (Supplementary Figures 5D and E). There was more information decoded from features of Mean, Median, Variance, and several multi-valued features, especially Wavelet and Orig Mag, compared to other features across the three datasets (Supplementary Figures 5A). The mentioned features dominated other features in terms of both average and maximum decoding accuracies (Supplementary Figures 5B and C). A complementary analysis suggested that there is a potential overlap between the neural codes that different features detected (Supplementary Figure 6).

We then directly compared of the Mean and the most informative variability-sensitive features (Wavelet and Orig Mag). Consistently across the datasets, there was moderate (3<BF<10) or strong (BF>10) evidence for higher decoding obtained by Wavelet and Orig Mag compared to the Mean feature on time points before 200 ms post-stimulus onset (Figure 3A). After 200 ms, this advantage sustained (Dataset 3), disappeared (Dataset 1) or turned into disadvantage (Dataset 2). Except for few very short continuous intervals, during which Wavelet provided higher decoding values compared to Orig Mag, the two features provided almost the same results (Figure 3; yellow dots on bottom panels). Comparing the parameters of the decoding curves, we found moderate (3<BF<10) or strong (BF>10) evidence for higher maximum decoding for the Wavelet and Orig Mag features than the Mean feature in Datasets 1 and 3 (Figure 3B). There was also moderate (3<BF<10) evidence for higher maximum decoding accuracy for Wavelet vs. Orig Mag (Figure 3B). There was also strong (BF>10) evidence for higher average decoding accuracy for the Wavelet and Orig Mag features over the Mean feature in Dataset 3 (Figure 3C). There was also moderate (3<BF<10) evidence for higher maximum decoding for Wavelet vs. Orig Mag in Datasets 2 and 3. These results show that the Wavelet feature provides the highest maximum (in Dataset 3) and average (in Datasets 2 and 3) decoding accuracies among the three features followed by the Orig Mag feature. The measures of maximum and average decoding accuracies were calculated in the post-stimulus onset (0-1000 ms) for each participant separately. We also compared the timing parameters of the decoding curves (i.e. the time to the first above-chance and maximum decoding relative to stimulus onset) obtained for the three features (Figure 3D and E), but found insufficient evidence (0.3<BF<3) for their difference.

Together, these results suggest that the inclusion of temporal variabilities of activity can provide additional information about object categories, to what is conventionally obtained from the Mean of activity. Note that, the advantage of Wavelet and Orig Mag features cannot be explained by the size/dimensionality of the feature space, as the number of dimensions were equalized across features. Importantly, however, the decoding of information from temporal variabilities did not lead to different temporal dynamics of information decoding. This can be explained by either the common cognitive processes producing the decoded neural codes (i.e. object categorization), the overlap between the information (neural codes) detected by our features or a combination of both.

### Do the features sensitive to temporal variabilities explain the behavioral recognition performance more accurately than the “mean” feature

Although we observed an advantage for the features which were sensitive to temporal variability (e.g. Wavelet) over other, more summarized, features (e.g. Mean), this can all be a by-product of more flexibility (e.g. inclusion of both ***temporal*** and ***spatial*** codes) in the former over the latter, and not read out by down-stream neurons that support behavior. To validate the behavioral relevance of the detected neural codes, we calculated the *correlation* between the decoding accuracies of features and the reaction times of participants (Vidaurre et al., 2019; Ritchie et al., 2015). Participants’ reaction times in object recognition have been previously shown to be predictable from decoding accuracy (Ritchie et al., 2015). We expected to observe negative correlations between the features’ decoding accuracies and participants’ reaction times in the post-stimulus span (Ritchie et al., 2015). This suggests that greater separability between neural representations of categories might lead to with categorizing them faster in behavior; supporting that the decoded neural codes might be used by neurons which drive behavior. We only used Dataset 2 in this analysis, as it was the only dataset with an active object detection task; therefore relevant reaction times were available. The (Spearman’s rank-order) correlations were calculated across the time course of the trials between the 10-dimensional vector of neural decoding accuracies obtained on every time point and the 10-dimensional vector of behavioral reaction times, both obtained from the group of 10 participants (Cichy et al., 2014). This resulted in a single correlation value for each time point for the whole group of participants.

All features, except Katz FD, showed negative trends after the stimulus onset (Figure 4A). The correlations showed more sustained negative values for the multi-valued vs. single-valued features (p<0.05). There was also larger negative peaks (generally < -0.5) for multi-valued features especially Wavelet, compared to other features (generally > -0.5). Specifically, while higher-order moment features (i.e. Variance, Skewness and Kurtosis) as well as many complexity features showed earlier negative peaks at around 150 ms, Mean, Median, frequency-domain features and multi-valued features showed later negative peaks after 300 ms. Therefore, the multi-valued features, especially Wavelet, which were sensitive to temporal variabilities of the signals, showed the most sustained and significant correlations to behavior.

**Figure 4.**
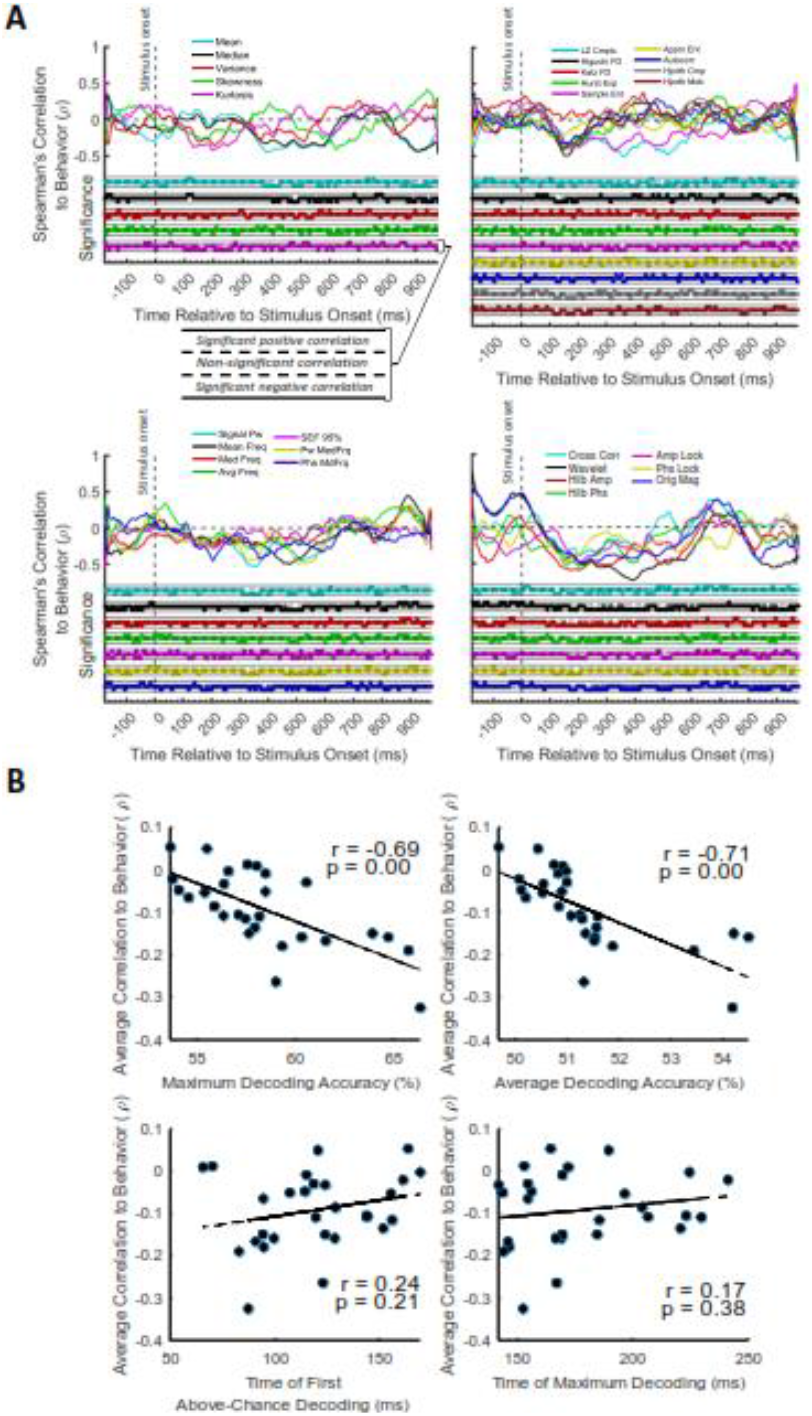
Correlation between the decoding accuracies and behavioral reaction times for Dataset 2 (other datasets did not have an active object recognition/detection task). (A) Top section in each panel shows the (Spearman’s) correlation coefficient obtained from correlating the decoding values and the reaction times for each feature separately. Correlation curves were obtained from the data of all participants. Bottom section shows positively or negatively significant (P<0.05; filled circles) or non-significant (p>0.05; empty circles) correlations as evaluated by random permutation of the variables in correlation. (B) Correlation between each of the amplitude and timing parameters of time-resolved decoding (i.e. maximum and average decoding accuracy and time of first and maximum decoding) with the average time-resolved correlations calculated from for the set of N=28 features. The slant line shows the best linear fit to the distribution of the data.

Visual inspection suggests that features which provided a higher decoding accuracy (e.g. Wavelet, Figure 3), did also better at predicting behavioral performance (e.g. Wavelet, Figure 4). To quantitatively see if such a relationship exists, we calculated the correlation between parameters of the decoding curves (introduced in Figure 3B-D) and the “average correlation to behavior” achieved by the same features (Figure 4A). Specifically, we used the “average” and “maximum” decoding accuracies, which we hypothesized to predict “average correlation to behavior”, and the “time of first above-chance” and “maximum” decoding accuracies (used as control variables here), which we hypothesized not to predict “average correlation to behavior”. To obtain the parameter of “average correlation to behavior”, we simply averaged the correlation to behavior in the post-stimulus time span for each feature separately (Figure 4A). Results showed that (Figure 4B), while the temporal parameters of “time of first above-chance” and “maximum” decoding (our control parameters) failed to predict the level of average correlation to behavior (r=0.24, p=0.21, and r=0.17, p=0.38, respectively), the parameters of “maximum” and “average” decoding accuracies significantly (r=-0.69 and r=-0.71 respectively, with p<0.0001; Pearson’s correlation) predicted the average correlation to behavior. Note the difference between the “Spearman’s correlation to behavior” calculated in Figure 4A and the correlations reported in Figure 4B. While the former is obtained by correlating the time-resolved decoding rates and corresponding reaction times across participants, the latter is calculated by correlating the post-stimulus average of the former correlations and their corresponding decoding parameters across features, rather than participants. This result suggests that the more effective the decoding of the neural codes, the better the prediction of behavior. Note that, this is not a trivial result; higher decoding values for the more informative features do not necessarily lead to higher correlation to behavior, as “correlation” normalizes the absolute values of input variables.

## Discussion

Temporal variability of neural activity has been suggested to provide an additional channel to the “mean” of activity for the encoding of several aspects of the input sensory information. This includes complexity (Garrett et al., 2020), uncertainty (Orbán et al., 2016) and variance (Hermundstad et al., 2014) of the input information. It is suggested that the brain optimizes the neuronal activation and variability to avoid over-activation (energy loss) for simple, familiar and less informative categories of sensory inputs. For example, face images, which have less variable compositional features, evoked less variable responses in fMRI, compared to house images, which were more varied, even in a passive viewing task (Garrett et al., 2020). This automatic and adaptive modulation of neural variability can result in more effective and accurate encoding of the sensory inputs in changing environments e.g. by suppressing uninformative neuronal activation for less varied (more familiar) stimuli such as face vs. house images (Garrett et al., 2020). Despite the recent evidence about the richness of information in temporal variability, which is modulated by the category of the sensory input (Garrett et al., 2020; Orbán et al., 2016; Waschke et al., 2021), majority of EEG studies still ignore variability in decoding. Specifically, they generally either extract variability (e.g. entropy and power) from the whole-trial activity (e.g. for brain-computer interface (BCI)) or use the simple “mean” (average) magnitude data within sub-windows of the trial (e.g. for time-resolved decoding; Grootswagers et al., 2017). The former can miss the informative within-trial variabilities/fluctuations of the trial in the highly dynamical and non-stationary evoked potentials. The latter, on the other hand, may overlook the informative variabilities within the sliding time windows as a result of temporal averaging.

Here, we quantified the advantage of the features sensitive to temporal variabilities over the conventional “mean” activity. In whole-trial analysis, we observed that, the features, which targeted informative sub-windows/samples of the trial (e.g. ERP components, Wavelet coefficients (Wavelet) and Original magnitude data (Orig Mag)), could provide more category information than the Mean feature, which ignored temporal variabilities. Interestingly, ERP components (N1, P2a and P2b) provided comparable results to that obtained by informative samples (Orig Mag) or Wavelet transformation (except for Dataset 3). That could be the reason for the remarkable decoding results achieved in previous studies which used ERPs (Wang et al., 2012; Qin et al., 2016) and Wavelet (Taghizadeh-Sarabi et al., 2015). These results also proposes that, we might not need to apply complex transformations (e.g. Wavelet) on the data in whole-trial analysis (Taghizadeh-Sarabi et al., 2015), as comparable results can be obtained using simple ERP components or original magnitude data. However, inclusion of more dimensions of the features in decoding or combining them (Karimi Rouzbahani et al., 2011; Qin et al., 2016) could potentially provide higher decoding accuracies for multi-valued (e.g. Wavelet; Taghizadeh-Sarabi et al., 2015) than ERP features (i.e. we equalized the dimensions across features here).

The Wavelet and Original magnitude data not only outperformed all the variability-sensitive features, but also the conventional Mean feature. Importantly, while features such as Hilbert phase and amplitude, Phase- and Amplitude-locking and Inter-electrode correlations, also had access to all the samples within the sliding analysis window, they failed to provide information comparable to Wavelet and Orig Mag features. The reason for the success of the Original magnitude data, seems to be that it basically makes no assumptions about the shape/pattern of the potential neural codes, as opposed to Hilbert phase (Hilb Phs), amplitude (Hilb Amp), and correlated variability (Cross Corr) each of which are sensitive ot one specific aspect of neural variability (i.e. phase, amplitude, correlation). The reason for success of the Wavelet feature, on the other hand, seems to be its reasonable balance between flexibility in detecting potential neural codes contained in the amplitude, phase and frequency/scale and a relatively lower susceptibility to noise as a result of filtering applied on different frequency bands (Guo et al., 2009). Together, these observations support the idea that neural codes are complex structures reflected in multiple aspects of EEG data e.g. amplitude, phase and frequency/scale (Panzeri et al., 2010; Waschke et al., 2021).

The advantage of Theta-over Broad-band in our data (Supplementary Figures 1 and 3) is consistent with previous monkey studies suggesting that Theta and Gamma frequency bands played major roles in feed-forward processing of visual information in the brain (Bastos et al., 2015), which also seemed dominant here (Supplementary Figure 4). One potential reason for the encoding of feed-forward information in the Theta band can be that bottom-up sensory signals transfer information about ongoing experiences, which might need to be stored in long-term memory for future use (Zheng and Colgin, 2015). Long-term memories are suggested to be encoded by enhanced long-lasting synaptic connections. The optimal patterns of activity which can cause such changes in synaptic weights were suggested to be successive Theta cycles which carry contents in fast Gamma rhythms (∼100 Hz; Larson et al., 1986). While direct correspondence between invasive vs. non-invasive neural data remains unclear (Ng et al., 2013), this study provides additional evidence for the major role of Theta frequency band in human visual perception (Wang et al., 2012; Qin et al., 2016; Jadidi et al., 2016; Taghizadeh-Sarabi et al., 2015; Torabi et al., 2017). It also suggests that BCI community might benefit from concentrating on specific frequency bands relevant to the cognitive or sensory processing undergoing in the brain; i.e. investigating the Theta band when stimulating the visual system.

One critical question for cognitive neuroscience has been whether (if at all) neuroimaging data can explain behavior (Williams et al., 2007; Ritchie et al., 2015; Woolgar et al., 2019; Karimi-Rouzbahani et al., 2019; Karimi-Rouzbahani et al., 2021a). We extended this question by asking whether more optimal decoding of object category information, can lead to better prediction of behavioral performance. We showed in our Dataset 2 that, this can be the case. Critically, here we observed for the same dataset that, there seems to be a linear relationship between the obtainable decoding accuracy and the explanatory power of the features. It implies that in order to bring neuroimaging observations closer to behavior, we might need to work on how we can read out the neural codes more effectively.

It has been suggested that neural variability is not only modulated by sensory information (as focused on here), but also by other top-down cognitive processes such as attention, expectation, memory and task demands (Waschke et al. 2021). For example, attention decreased low-frequency neural variabilities/power (2-10 Hz; which is referred to as “desynchronization”) while increasing high-frequency neural variabilities/power (Wyart and Tallon-Baudry, 2009). Therefore, in the future, it will be interesting to know which features best detect the modulation of neural variability in other cognitive tasks. Moreover, it is interesting to know how (if at all) a combination of the features used in this study could provide any additional information about object categories and/or behavior. In other words, although all of the individual features evaluated here covered some variance of category object information, to detect the neural information more effectively, it might be helpful to combine multiple features using supervised and un-supervised methods (Karimi Rouzbahani et al., 2011; Qin et al., 2016).

The cross-dataset, large-scale analysis methods implemented in this study aligns with the growing trend towards meta-analysis in cognitive neuroscience. Recent studies have also adopted and compared several datasets to facilitate forming more rigorous conclusions about how the brain performs different cognitive processes such as sustained attention (Langner et al., 2013) or working memory (Adam et al., 2020). Our results provide evidence supporting the idea that neural variability seems to be an additional channel for information encoding in EEG, which should not be simply ignored.

## Acknowledgements

This research was funded by the Royal Society’s Newton International Fellowship SUAI/059/G101116 to Hamid Karimi-Rouzbahani.

## Supplementary Materials

**Supplementary Figure 1.**
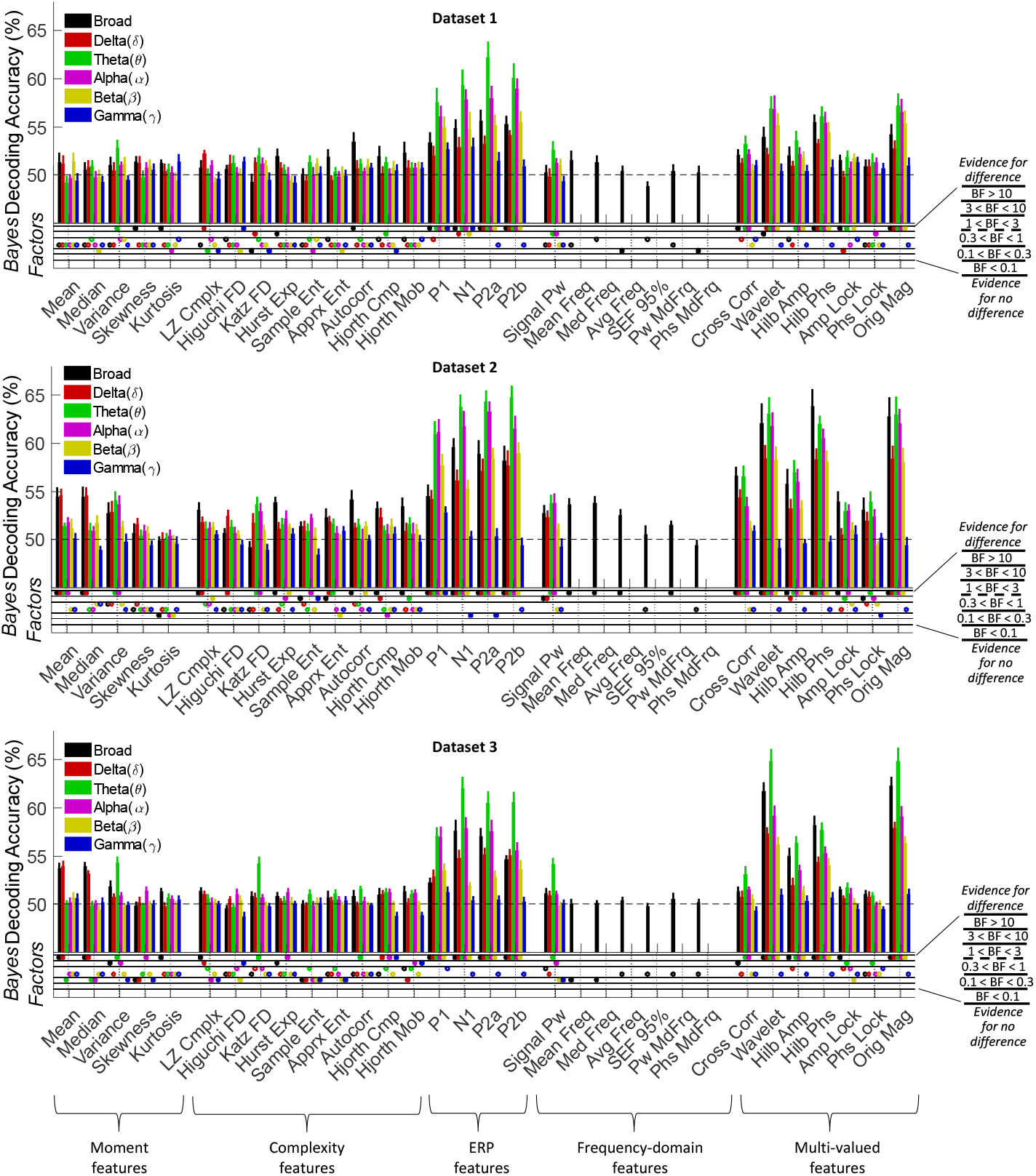
Whole-trial decoding of object categories from the three datasets using 32 features in different frequency bands (for Bayesian evidence analyses see Supplementary Figure 2). Decoding of category information using the 32 features in the 6 frequency bands. The black horizontal dashed lines on the top panel refer to chance-level decoding. Thick bars show the average decoding across participants (error bars Standard Error across participants). Bayes Factors are shown in the bottom panel of each graph: Filled circles show moderate/strong evidence for either hypothesis and empty circles indicate insufficient evidence. They show the results of Bayes factor analysis when evaluating the difference from chance-level decoding.

**Supplementary Figure 2.**
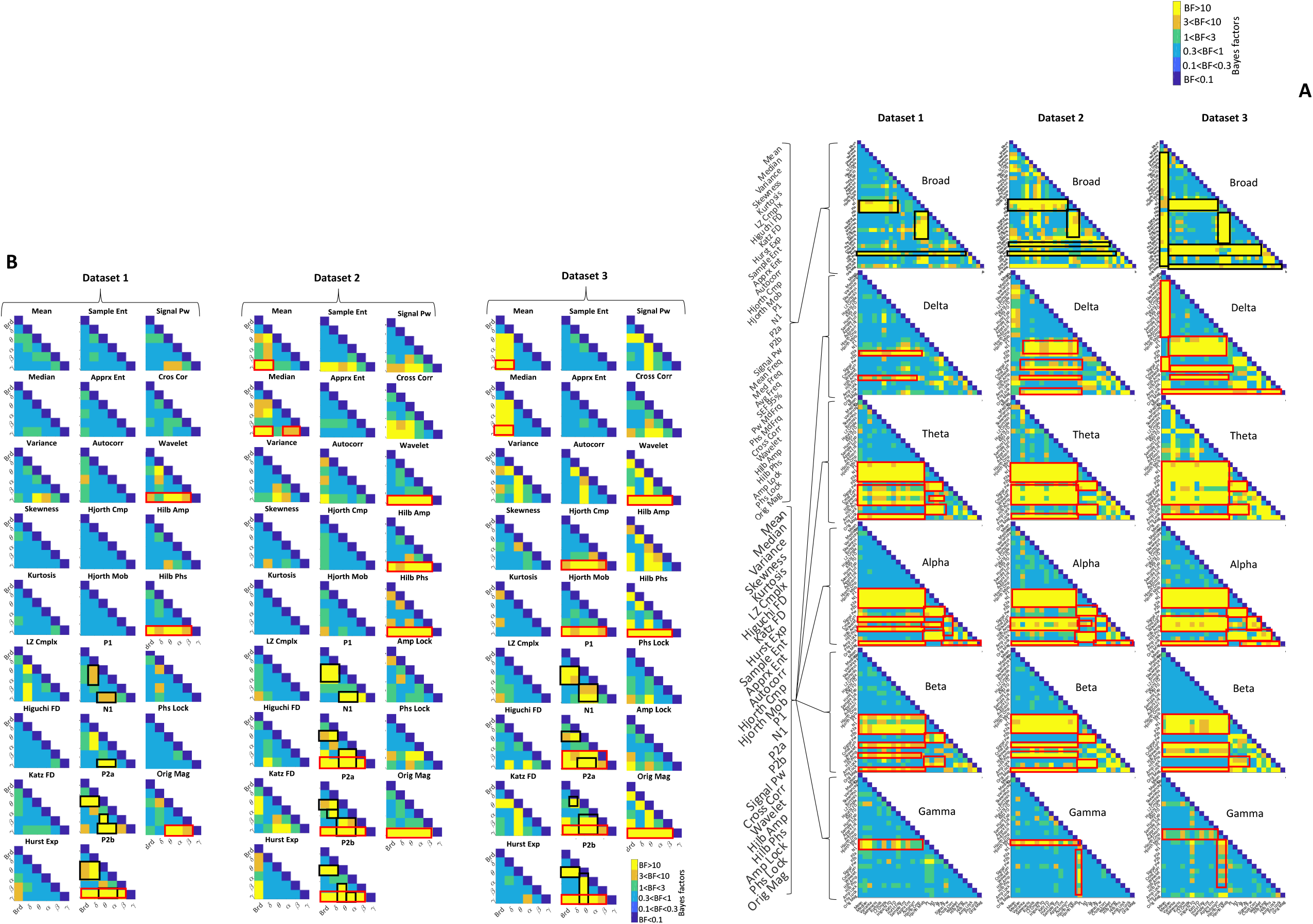
(A) Bayes factor matrices comparing whole-trial decoding results across different frequency bands and dataset separately. Matrices show different levels of evidence for existing difference (moderate 3<BF<10, Orange; strong BF>10, Yellow), no difference (moderate 0.1<BF<0.3, light blue; strong BF<0.1, dark blue) or insufficient evidence (1<BF<3 green; 0.3<BF<1 Cyan) for either hypotheses. Black and red boxes indicate moderate or strong evidence for higher decoding values for specific features mentioned and compared in the text. For example, for Dataset 1, there is insufficient evidence for difference between decoding values of most features in the Gamma band as indicated by the light blue color in most cells. However, there is moderate or strong evidence that Mean and Median features are different from N1 and P1 as indicated by yellow color and the decoding accuracies in Supplementary Figure 1. (B) Bayes factor matrices comparing whole-trial decoding results within each frequency band, across features separated by datasets. Black and red boxes indicate moderate or strong evidence for higher decoding values for specific features mentioned and compared in the text.

**Supplementary Figure 3.**
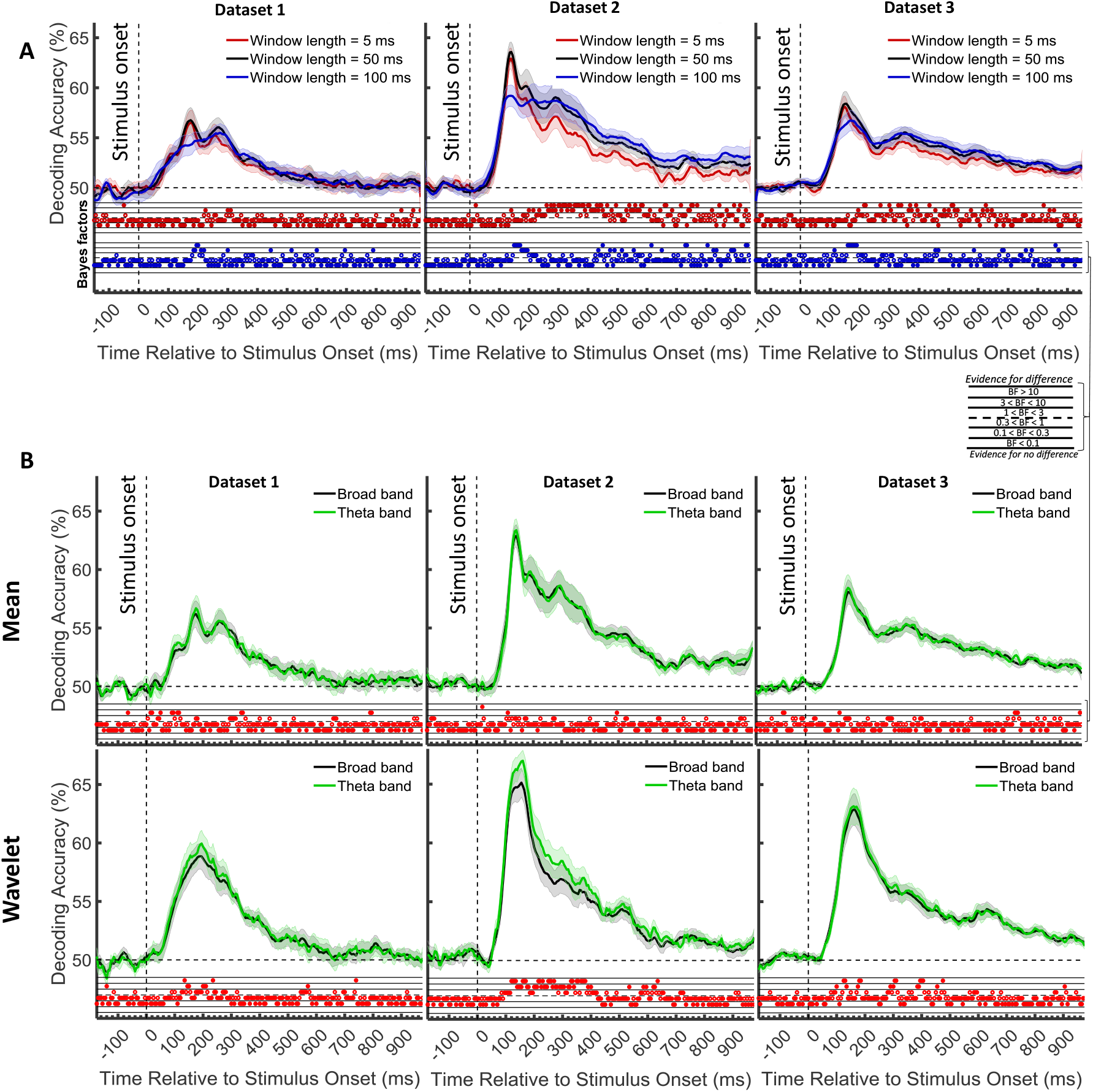
We selected the window length of 50 ms for our time-resolved analyses because it was neither too long to hide the true temporal dynamics of information processing in the brain, nor too short to avoid the proper calculation of features (e.g. complexity and multi-valued). To assure that we did not miss the true obtainable dynamic range (amplitude) of accuracies, we compared category decoding obtained from time windows of 5 (i.e. which was the case in most previous studies all of which relied on signals’ mean (Grootswagers et al., 2017; Karimi-Rouzbahani et al., 2017b) and 100 ms with that used here from 50 ms time windows. Consistently across the three datasets, results showed that the highest decoding accuracies were obtained from the 50 ms time windows, both in terms of maximum and average decoding accuracy after the stimulus onset. Interestingly, lengthening the time windows decreased the maximum decoding but increased the decoding accuracies in the later stages of the processing (i.e. from 200 ms onwards; probably after initial hard-wired processing of visual stimuli). This may suggest that later stages of category processing (probably involving feedback/recurrent processing; which are activated by the longer presentation time in datasets 2 and 3), take longer processing times, therefore captured better using longer time windows. (A) Comparison of decoding accuracies using different length for the sliding time window. The bottom section shows the Bayes factor evidence for the difference between the 50 ms window and the other two window lengths. (B) Comparison of decoding accuracies using different frequency bands for the Mean (top) and Wavelet (bottom) features. Each column shows the results for one dataset. Top section in each panel shows the decoding accuracies across time and the horizontal dashed lines on the top panel refer to chance-level decoding. Filled circles in the Bayes Factors show moderate/strong evidence for either difference or no difference between the decoding curves and empty circles indicate insufficient evidence for either hypotheses. Thick lines show the average decoding accuracy across participants (error bars Standard Error across participants).

**Supplementary Figure 4.**
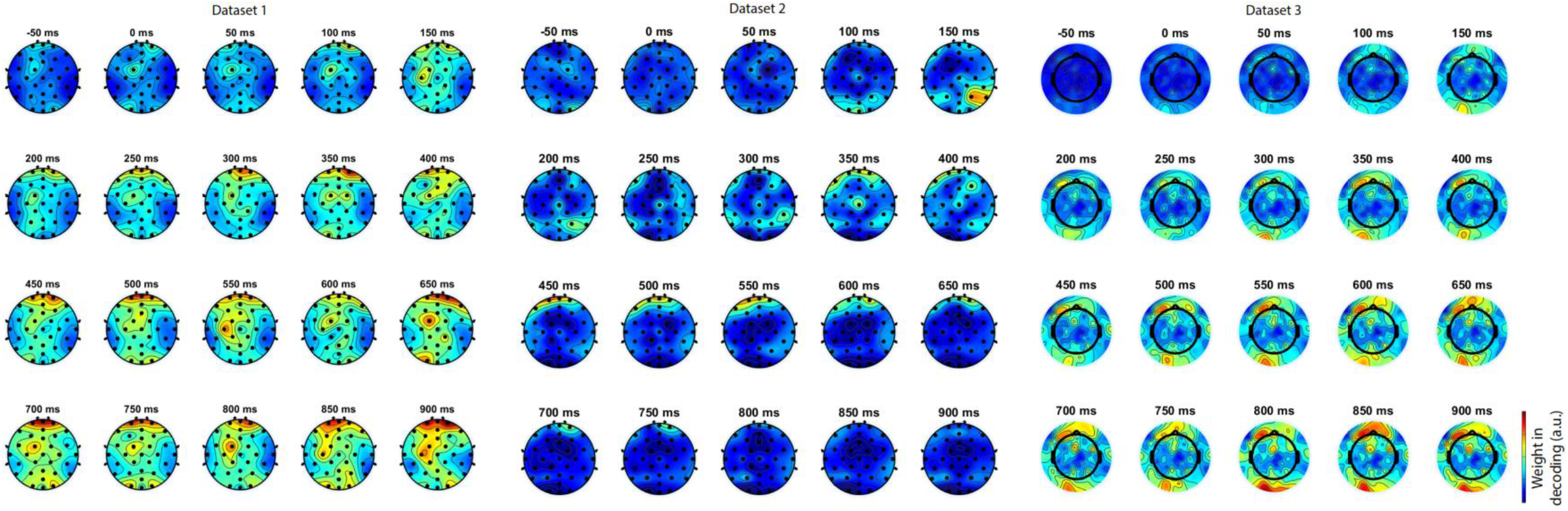
To see if that the Theta frequency band supports feed-forward flow of information in our datasets, we also calculated spatial maps of classifier weights on the head for the conventionally used Mean feature in the Theta frequency band. These classifier weights reflect how much information each electrode provides about object categories at different time points. The categorical object information initially appeared ∼50 or ∼100 ms after the stimulus onset in all three datasets predominantly in the occipital areas. This was followed in later time windows (∼100 ms to ∼150 ms) by the information appearing in both the occipital (all datasets), occipito-temporal (all datasets), central (Dataset 1) as well as frontal electrodes (all datasets). Finally, from around ∼300 ms onwards, the object category information seemed to be dominantly represented in occipital and frontal (Datasets 1 and 3) areas or only the frontal (Dataset 2) area. These results seem to support feed-forward flow of information through the ventral and dorsal visual streams as well as from occipital to frontal brain areas during the trial. However, based on the limited spatial resolution of EEG and the susceptibility of classifier weights to artefacts (Haufe et al., 2014), we should be careful not to over-interpret these spatiotemporal maps. Classifier weights in decoding. These topographic classifier maps were obtained from classifier weight values provided by the LDA classifiers used in decoding. The weight values have different scales for different datasets based on the nature of the data. Therefore, we normalized them for presentation within each dataset for clearer presentation. Hot colors show higher and cold colors reflect lower weights.

**Supplementary Figure 5.**
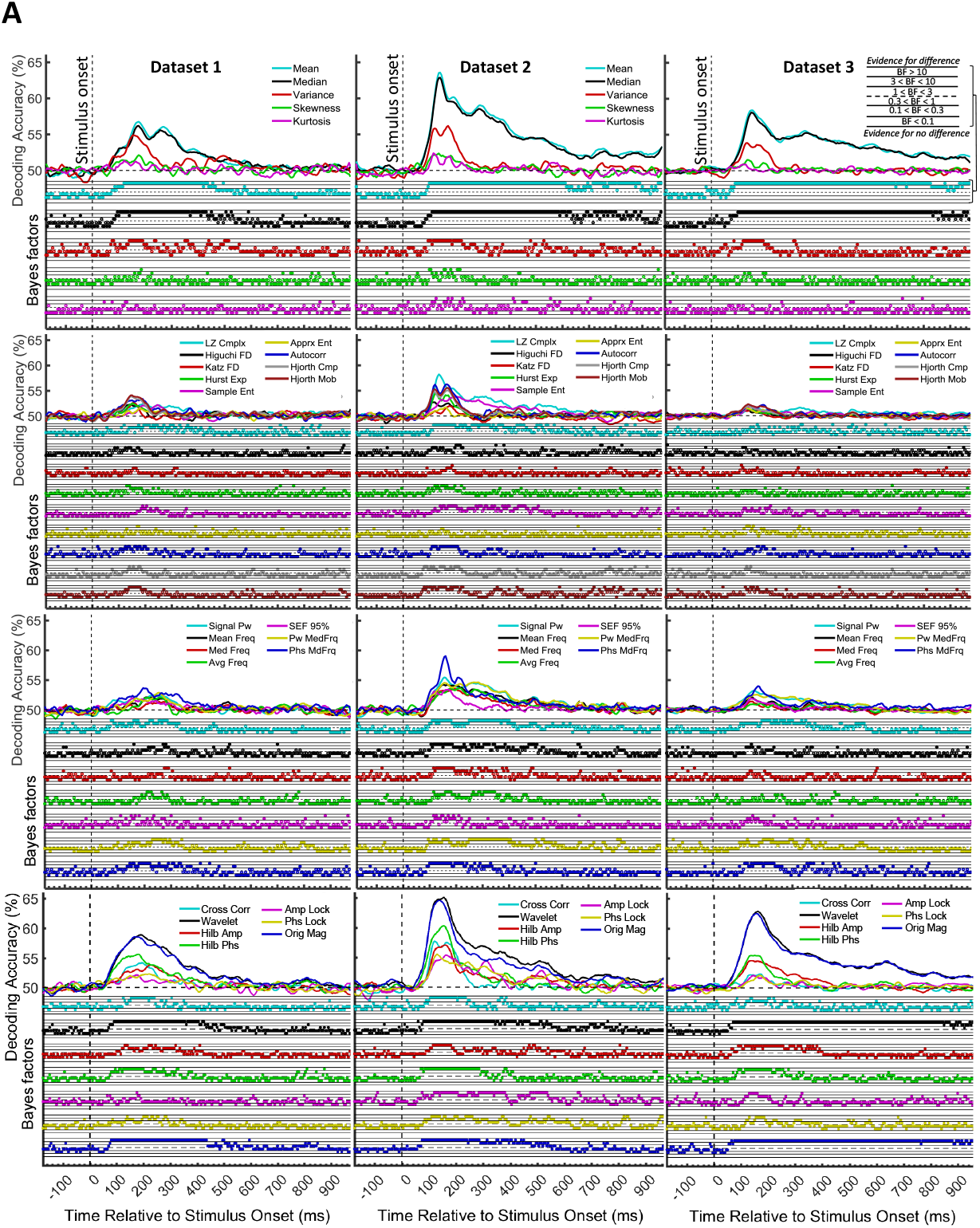
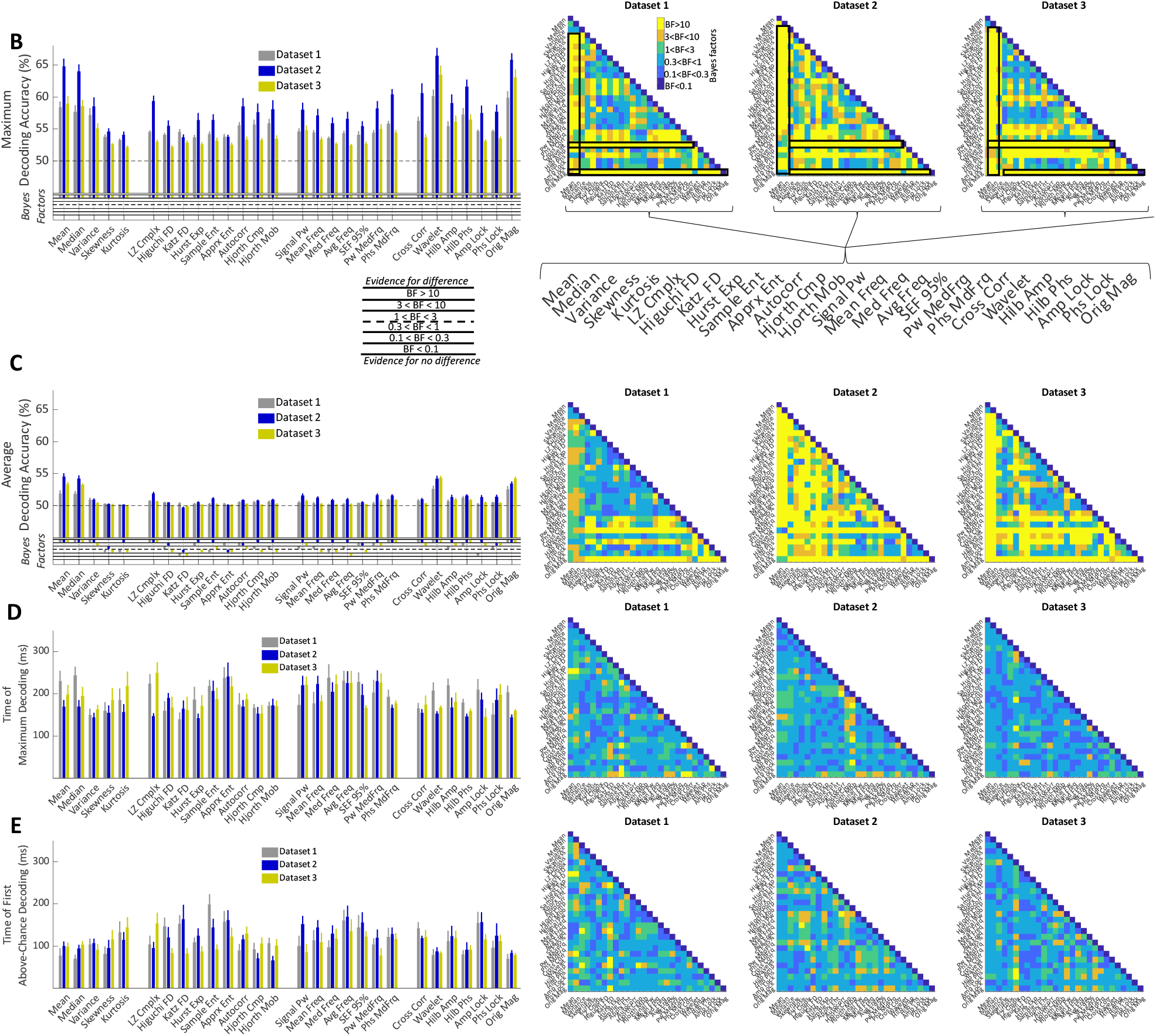
Time-resolved decoding of object categories from the three datasets using 28 features and Bayesian evidence analyses. Each row shows the results of one type of feature (i.e. moment, complexity, frequency-domain and multi-valued features from top to bottom, respectively). Curves show the average decoding across participants. Each column shows the results for one dataset. Top section in each panel shows the decoding accuracies across time and the bottom section shows the Bayes factor evidence for the difference of the decoding accuracy compared to chance-level decoding. The horizontal dashed lines on the top panel refer to chance-level decoding. Filled circles in the Bayes Factors show moderate/strong evidence for either difference or no difference from chance-level decoding and empty circles indicate insufficient evidence for either hypotheses. Timing and amplitude parameters extracted from the time-resolved accuracies of each feature and each dataset and their Bayesian evidence analyses. (B-E) Left: the maximum and average decoding accuracies, the time of maximum and the first above-chance decoding. Thick bars show the average across participants (error bars Standard Error across participants). Bottom section on B and C show the Bayes factor evidence for the difference of the decoding accuracy compared to chance-level decoding; Right: matrices compare the right parameters obtained from different features. Different levels of evidence for existing difference (moderate 3<BF<10, Orange; strong BF>10, Yellow), no difference (moderate 0.1<BF<0.3, light blue; strong BF<0.1, dark blue) or insufficient evidence (1<BF<3 green; 0.3<BF<1 Cyan) for either hypotheses. Black and red boxes show moderate or strong evidence for higher decoding values for specific features compared other sets of features as explained in the text. The horizontal dashed lines on the left panels of (B) and (D) refer to change-level decoding. Filled circles in the Bayes Factors show moderate/strong evidence for either hypothesis and empty circles indicate insufficient evidence.

**Supplementary Figure 6.**
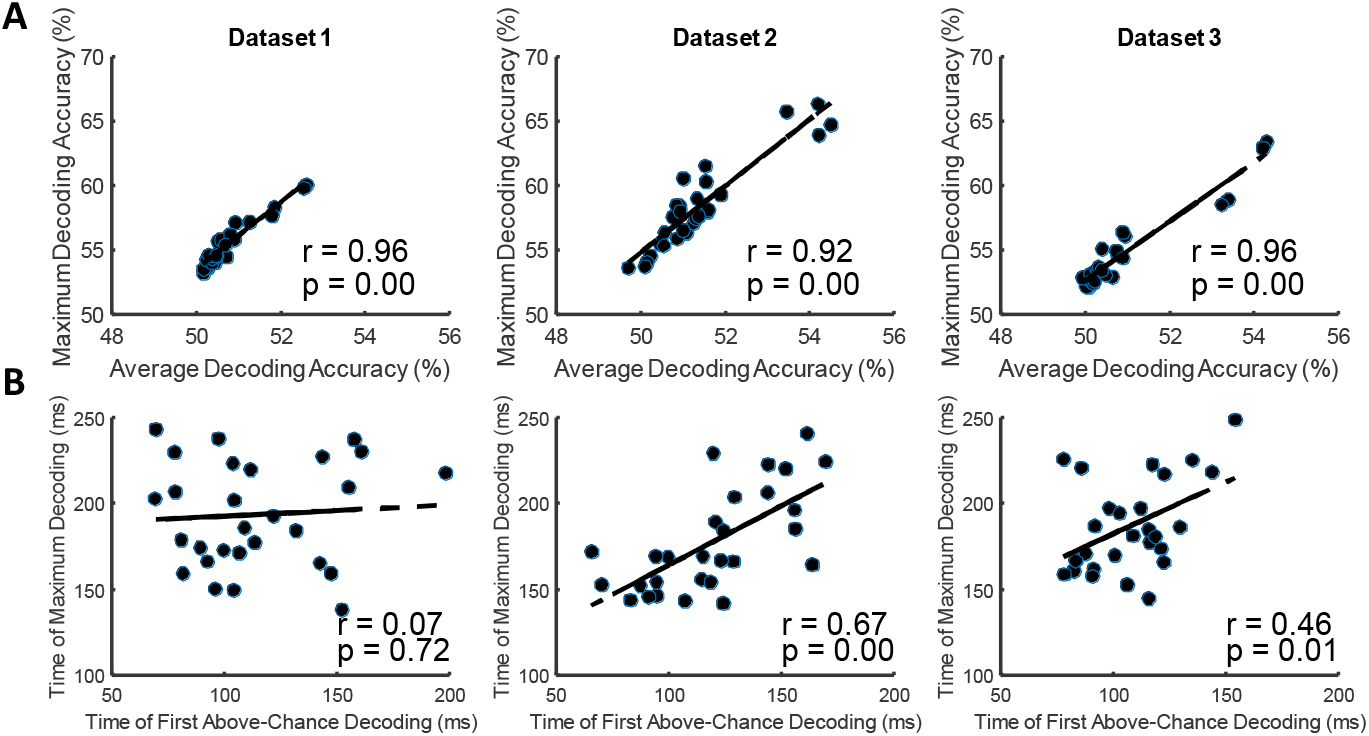
The temporal dynamics of different features seem to reflect a similar decoding pattern in the sense that the most informative features can lead to both a higher maximum decoding and a more sustained decoding pattern along the trial and vice versa. This suggests that there might be a general advantage for the more vs. less informative features which is reflected both in their maxima as well as their sustained decoding patterns. Alternatively, it can be the case that there is no relationship between the maxima and the average decoding across features, suggesting that each feature might detect different neural codes. To test this question, we calculated the correlation between the average and maximum decoding values for all features, which showed highly correlated results (r > 0.9; p < 0.01; Supplementary Figure 6A). This suggests that, all features followed a generally similar pattern of decoding with more informative features providing higher decoding maxima and a more sustained level of information decoding. There has been no consensus yet about whether the time of the maximum or the first above-chance decoding reflects the speed of category processing in the brain (Grootswagers et al., 2017; Ritchie et al., 2015). Hence, we calculated the correlation of these temporal parameters across features to see if they both possibly reflect the dynamics of the same processing mechanism in the brain. The time of first above-chance and maximum decoding correlated in Datasets 2 and 3 but not Dataset 1 (r=0.67, r=0.51 and r=0.07 respectively for Datasets 1, 2 and 3; Supplementary Figure 6B). Lack of significant correlation for Dataset 1 can be explained by the lower decoding values in Dataset 1 compared to the other datasets making the correlations noisier. Therefore, features that reached their above-chance decoding earlier also reached their maximum decoding earlier leading to the suggestion that they both reflect the temporal dynamics of the same cognitive processes with some delay. Correlation between the pairs of amplitude (A) and timing (B) parameters of the time-resolved decoding (i.e. maximum and average decoding accuracy and time of first and maximum decoding) for the set of N=28 individual features. The slant line shows the best linear fit to the distribution of the correlation data.

## Footnotes

1 https://purl.stanford.edu/tc919dd5388

2 https://www.mathworks.com/matlabcentral/fileexchange/38211-calc_lz_complexity

3 https://ww2.mathworks.cn/matlabcentral/fileexchange/50290-higuchi-and-katz-fractal-dimension-measures

4 https://www.mathworks.com/matlabcentral/fileexchange/9842-hurst-exponent

5 https://www.mathworks.com/matlabcentral/fileexchange/32427-fast-approximate-entropy

